# Stepwise activation mechanism of the scramblase nhTMEM16 revealed by cryo-EM

**DOI:** 10.1101/455287

**Authors:** Valeria Kalienkova, Vanessa Clerico Mosina, Laura Bryner, Gert T. Oostergetel, Raimund Dutzler, Cristina Paulino

**Affiliations:** Department of Biochemistry, University of Zürich, Winterthurer Str. 190, CH-8057 Zürich, Switzerland; Department of Structural Biology at the Groningen Biomolecular Sciences and Biotechnology Institute, University of Groningen, Nijenborgh 7, 9747 AG Groningen, The Netherlands

## Abstract

Scramblases catalyze the movement of lipids between both leaflets of a bilayer. Whereas the X-ray structure of the protein nhTMEM16 has previously revealed the architecture of a Ca^2+^-dependent lipid scramblase, its regulation mechanism has remained elusive. Here, we have used cryo-electron microscopy and functional assays to address this question. Ca^2+^-bound and Ca^2+^-free conformations of nhTMEM16 in detergent and lipid nanodiscs illustrate the interactions with its environment and they reveal the conformational changes underlying its activation. In this process, Ca^2+^-binding induces a stepwise transition of the catalytic subunit cavity, converting a closed cavity that is shielded from the membrane in the absence of ligand, into a polar furrow that becomes accessible to lipid headgroups in the Ca^2+^-bound state. Additionally, our structures demonstrate how nhTMEM16 distorts the membrane at both entrances of the subunit cavity, thereby decreasing the energy barrier for lipid movement.

**Impact statement:** cryo-EM reveals the properties of distinct conformations occupied during activation of the lipid scramblase nhTMEM16 and provides new insights into its interactions with the lipid environment.

## Introduction

The movement of lipids between both leaflets of a phospholipid bilayer, which is also referred to as lipid flip-flop, is energetically unfavorable since it requires the transfer of the polar headgroup from its aqueous environment across the hydrophobic core of the membrane (Kornberg and McConnell, 1971). Consequently, spontaneous lipid flip-flop is rare and occurs at a time scale of several hours. Whereas the sparsity of this event permits the maintenance of lipid asymmetry, there are processes that require the rapid transbilayer movement of lipids (Bevers and Williamson, 2016; Sanyal and Menon, 2009). Proteins that facilitate this movement by lowering the associated high energy barrier are termed lipid scramblases (Pomorski and Menon, 2006; Williamson, 2015). Lipid scramblases are generally non-selective and do not require the input of energy. They participate in various cellular functions ranging from lipid signaling during blood coagulation and apoptosis to the synthesis of membranes, cell division and exocytosis (Nagata et al., 2016; Whitlock and Hartzell, 2016). First structural insight into lipid scrambling was provided from the X-ray structure of the fungal protein from *Nectria haematococca*, termed nhTMEM16 (Brunner et al., 2014). This protein is part of the TMEM16 family (Milenkovic et al., 2010), which encompasses calcium-activated ion channels and lipid scramblases with a conserved molecular architecture (Brunner et al., 2016; Caputo et al., 2008; Falzone et al., 2018; Schroeder et al., 2008; Suzuki et al., 2013; Suzuki et al., 2010; Yang et al., 2008). As common for the family, nhTMEM16 is a homodimer with subunits containing a cytoplasmic domain and a transmembrane unit composed of 10 membrane-spanning segments. The lipid permeation path is defined by a structural entity termed the ‘subunit cavity’, which is located at the periphery of each subunit. In the X-ray structure, which was obtained in the presence of Ca^2+^, this site exposes a continuous polar cavity that spans the entire membrane and that is of appropriate size to harbor a lipid headgroup (Brunner et al., 2014).

The ‘subunit cavity’ is believed to provide a pathway for the polar moieties of lipids across the membrane, whereas the apolar acyl chains remain embedded in the hydrophobic core of the membrane (Bethel and Grabe, 2016; Brunner et al., 2014; Jiang et al., 2017; Lee et al., 2018; Malvezzi et al., 2018; Stansfeld et al., 2015). This process resembles a credit-card mechanism for lipid scrambling, which was previously proposed based on theoretical considerations (Pomorski and Menon, 2006). Although in this structure, Ca^2+^ ions were identified to bind to a conserved site in the transmembrane domain, the mechanism of how Ca^2+^ activates the protein and how the protein interacts with the surrounding lipids remained elusive. To gain insight into these questions, we have determined structures of the protein in Ca^2+^-bound and Ca^2+^-free conformations by single particle cryo-electron microscopy (cryo-EM), both in detergent and in a membrane environment and we have characterized the effect of mutants on the activation process. Our data reveal two essential features of the protein. They show how the protein deforms the membrane to lower the energy barrier for lipid movement and to steer translocating lipids towards the subunit cavity. Additionally, they define the conformational changes leading to the opening of the subunit cavity, which is shielded from the membrane in a Ca^2+^-free conformation, but exposes a hydrophilic furrow for the permeation of lipids upon Ca^2+^ binding in a stepwise process involving coupled ligand-dependent and ligand-independent steps.

## Results

### Structural characterization of nhTMEM16 in detergent

For our investigations of the ligand-induced activation mechanism of the lipid scramblase nhTMEM16, we have recombinantly expressed the protein in the yeast *S. cerevisiae*, purified it in the detergent n-Dodecyl β-D-maltoside (DDM) in the absence of Ca^2+^ and added Ca^2+^ during sample preparation for structure determination and transport experiments. We have reconstituted the purified protein into liposomes of different composition containing fluorescently labeled lipids and characterized lipid transport using an assay that follows the irreversible decay of the fluorescence upon addition of the membrane-impermeable reducing agent dithionite to the outside of the liposomes (Brunner et al., 2014; Malvezzi et al., 2013; Ploier and Menon, 2016). In these experiments we find a Ca^2+^-dependent increase of transport but also a pronounced basal activity of the protein in absence of Ca^2+^, which was previously described as a hallmark of fungal TMEM16 scramblases (Brunner et al., 2014; Malvezzi et al., 2013) (Figure 1-figure supplement 1). For the structural characterization of nhTMEM16, we have initially studied Ca^2+^-bound and Ca^2+^-free samples in detergent by cryo-EM. By this approach, we have obtained two datasets of high quality and of sufficient resolution (*i.e.* 3.6 Å for Ca^2+^-bound and 3.7 Å for Ca^2+^-free conditions) for a detailed interpretation by an atomic model (Figure 1; Figure 1-figure supplements 2-4; Table 1). In the structure determined in the presence of Ca^2+^, we find a conformation of the protein, which closely resembles the X-ray structure with a root mean square deviation (RMSD) of Cα atoms of 0.66 Å between the two structures (Figure 1D,E). The high similarity prevails irrespective of the absence of crystal packing interactions and despite of the fact that the sample used for cryo-EM was only briefly exposed to Ca^2+^ at a comparably low (*i.e.* 300 µM) concentration, whereas for the X-ray structure, the protein was exposed to Ca^2+^ during purification and crystallization at a 10-fold higher concentration (Brunner et al., 2014). Cryo-EM density at the conserved Ca^2+^-binding site, contained within each subunit of the homodimer, clearly indicates the presence of two bound Ca^2+^ ions located at the position originally identified in the X-ray structure (Figure 1F) (Brunner et al., 2014). Remarkably, we found a very similar structure of the protein in the absence of Ca^2+^, thus underlining the stability of this conformation in detergent solution (Figure 1G,H). Although both datasets display equivalent conformations of the protein, the structure determined in the absence of Ca^2+^ clearly shows an altered distribution of the divalent ligand in its binding site, concomitant with a weaker density of the C-terminal part of α6 indicating its increased mobility (Figure 1G,I; Figure1-figure supplement 3). The same region was previously identified as key structural element in the Ca^2+^-regulation of TMEM16A (Dang et al., 2017; Paulino et al., 2017a; Peters et al., 2018). In the cryo-EM density, both strong peaks of the ligand found in the Ca^2+^-bound conformation are absent and instead replaced by weak residual density at low contour located at the position of the lower Ca^2+^ ion (Figure 1F,I). Due to the presence of millimolar amounts of EGTA in the buffer, the resulting free Ca^2+^-concentration is in the pM range, well below the estimated nM potency of the divalent cation (Figure 1-figure supplement 1). Thus, whereas we do not want to exclude the possibility of Ca^2+^ binding at low occupancy, this density might as well correspond to a bound Na^+^ ion, which is present in the sample buffer at a concentration of 150 mM (*i.e.* at a 10^10^ times higher concentration compared to Ca^2+^). We think that the stability of the open conformation displayed in the structures of the protein in detergent, irrespectively of the presence of bound Ca^2+^, does not reflect an intrinsic rigidity of the subunit cavity. Instead, it is likely a consequence of experimental conditions, which stabilize the observed conformation of the protein. This is in accordance with residual density from the headgroup of a bound detergent molecule at the upper part of the lipid translocation path found in both datasets, which might prevent subunit cavity closure (Figure 1-figure supplement 5). Interestingly, the detergent molecule is interacting with Tyr 439, a residue that was shown to be essential for lipid translocation in MD simulations and functional studies (Lee et al., 2018). The Ca^2+^-free structure in detergent thus likely displays a conformation that is responsible for the basal activity observed for nhTMEM16 and other fungal scramblases (Figure 1-figure supplement 1) (Brunner et al., 2014; Malvezzi et al., 2013). We expect this state to be less populated in a lipid bilayer.

**Table 1.** Cryo-EM data collection, refinement and validation statistics.

|  | nhTMEM16,<br>DDM, +Ca<sup>2+</sup><br>(EMDB-4588,<br>PDB 6QM5) | nhTMEM16,<br>DDM, -Ca<sup>2+</sup><br>(EMDB-4589,<br>PDB 6QM6) | nhTMEM16,<br>2N2, -Ca<sup>2+</sup><br>(EMDB-4587,<br>PDB 6QM4) |
| --- | --- | --- | --- |
| **Data collection and processing** |  |  |  |
| Microscope | FEI Talos Arctica | FEI Talos Arctica | FEI Talos Arctica |
| Camera | Gatan K2 Summit + | Gatan K2 Summit + | Gatan K2 Summit + |
| Magnification | GIF | GIF | GIF |
|  | 49,407 | 49,407 | 49,407 |
| Voltage (kV) | 200 | 200 | 200 |
| Exposure time frame/total (s) | 0.15/9 | 0.15/9 | 0.15/9 |
| Number of frames per image | 60 | 60 | 60 |
| Electron exposure (e-/Å<sup>2</sup>) | 52 | 52 | 52 |
| Defocus range (μm) | -0.5 to -2.0 | -0.5 to -2.0 | -0.5 to -2.0 |
| Pixel size (Å) | 1.012 | 1.012 | 1.012 |
| Box size (pixels) | 220 | 240 | 240 |
| Symmetry imposed | C2 | C2 | C2 |
| Initial particle images (no.) | 251,693 | 570,203 | 1,379,187 |
| Final particle images (no.) | 120,086 | 238,070 | 133,961 |
| Map resolution (Å) |  |  |  |
| 0.143 FSC threshold | 3.64 | 3.68 | 3.79 |
| Map resolution range (Å) | 3.4-5.0 | 3.4-5.0 | 3.3-5.0 |
| **Refinement** |  |  |  |
| Initial model used | PDB 4WIS | 6QM5 | 6QMB |
| Model resolution (Å) | 3.6 | 3.7 | 3.8 |
| FSC threshold |  |  |  |
| Model resolution range (Å) | 80-3.6 | 80-3.7 | 80-3.8 |
| Map sharpening *B* factor (Å<sup>2</sup>) | -126 | -147 | -150 |
| Model composition |  |  |  |
| Nonhydrogen atoms | 10874 | 10852 | 10578 |
| Protein residues | 1346 | 1344 | 1308 |
| Ligands | 4 | - | - |
| *B* factors (Å<sup>2</sup>) |  |  |  |
| Protein | 55.75 | 76.42 | 94.09 |
| Ligand | 33.72 | - | - |
| R.m.s. deviations |  |  |  |
| Bond lengths (Å) | 0.007 | 0.004 | 0.007 |
| Bond angles (°) | 0.821 | 0.835 | 0.983 |
| Validation |  |  |  |
| MolProbity score | 1.47 | 1.42 | 1.53 |
| Clashscore | 4.02 | 4.35 | 4.09 |
| Poor rotamers (%) | 0 | 0 | 0 |
| Ramachandran plot |  |  |  |
| Favored (%) | 95.92 | 96.74 | 95.17 |
| Allowed (%) | 4.08 | 3.26 | 4.83 |
| Disallowed (%) | 0 | 0 | 0 |

**Figure 1.**
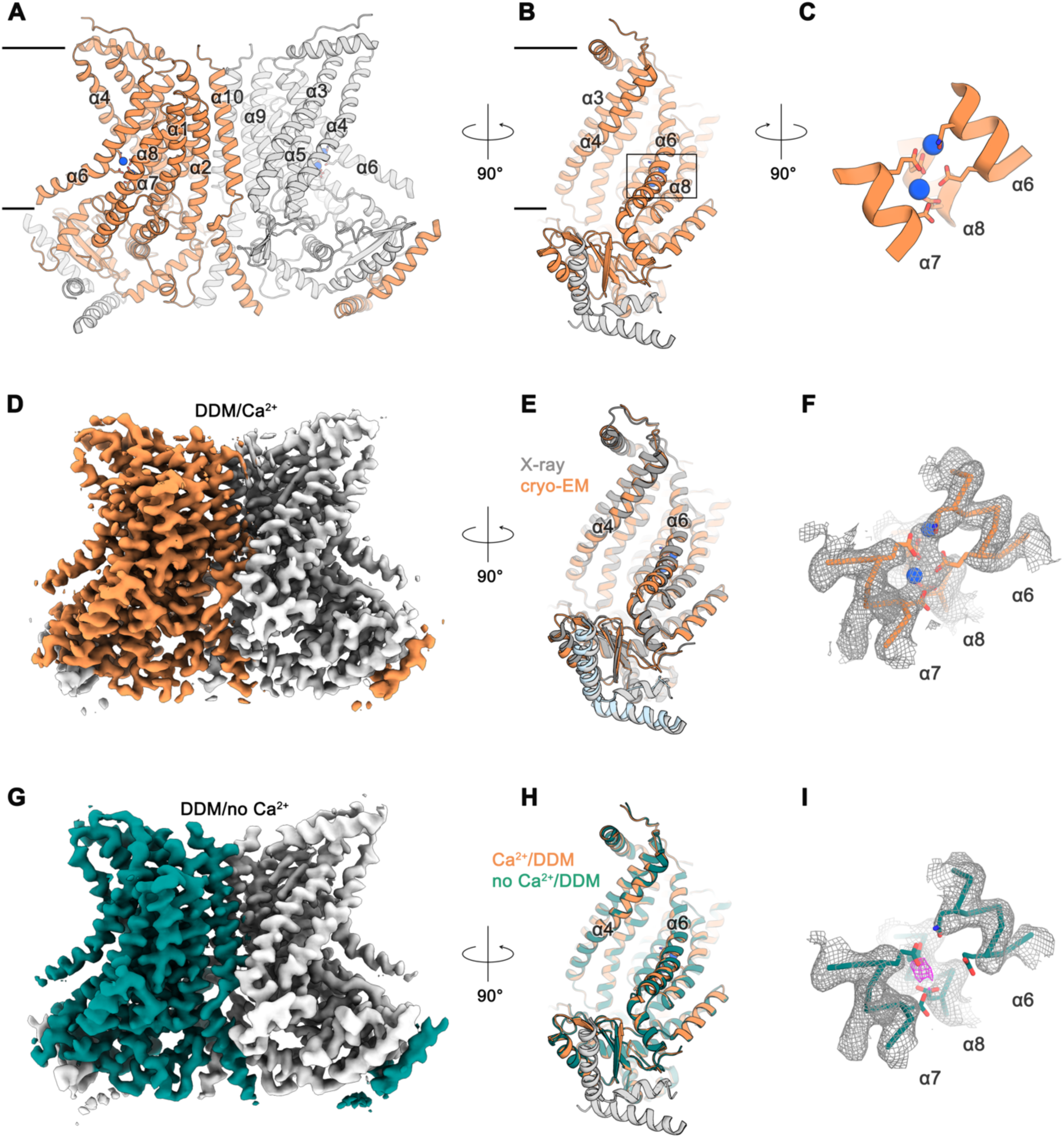
Cryo-EM structures of nhTMEM16 in detergent. Ribbon representation of the Ca^2+^-bound cryo-EM structure of nhTMEM16 in detergent. The view is from within the membrane perpendicular to the long dimension of the protein (**A**), towards the subunit cavity (**B**) and at the Ca^2+^-ion binding site (**C**). The membrane boundary is indicated. Subunits in the dimer are depicted in orange and grey, respective transmembrane-helices are labelled and Ca^2+^-ions are displayed as blue spheres. Relative orientations are indicated and the location of the Ca^2+^-binding site is highlighted by a box in panel B. (**D**) Cryo-EM map of the nhTMEM16 dimer (orange and grey) in DDM in presence of Ca^2+^ at 3.6 Å sharpened with a b-factor of –126 Å^2^ and contoured at 6 σ. (**E**) Ribbon representation of a superposition of the Ca^2+^-bound structure in detergent determined by cryo-EM (orange and grey) and the Ca^2+^-bound X-ray structure (dark grey and light blue, PDBID: 4WIS) The view is as in panel B. (**F**) View of the Ca^2+^-ion binding site of the Ca^2+^-bound state of nhTMEM16 in DDM. Ca^2+^-ions are displayed as blue spheres. (**G**) Cryo-EM map of the nhTMEM16 dimer (green and grey) in DDM in the absence of Ca^2+^ at 3.7 Å sharpened with a b-factor of –147 Å^2^ and contoured at 6 σ. (**H**) Ribbon representation of a superposition the Ca^2+^-bound (orange and grey) and the Ca^2+^-free structure (green and grey) in detergent determined by cryo-EM. The view is as in panel B. (**I**) View of the Ca^2+^-binding site in the Ca^2+^-free state of nhTMEM16 in DDM. The location of weak residual density at the center of the Ca^2+^-binding site is indicated in magenta. F,I The respective cryo-EM densities are contoured at 7 σ and shown as mesh. The backbone is displayed as Cα-trace, selected side-chains as sticks.

### Structure of the Ca^2+^-free state of nhTMEM16 in lipid nanodiscs

The absence of conformational changes in the subunit cavity of nhTMEM16 in detergents motivated us to study the structure of the protein in a lipid bilayer. For that purpose, we have reconstituted nhTMEM16 into nanodiscs of different sizes using a lipid composition at which the protein retains its lipid scrambling activity, although with considerably slower kinetics (Figure 1-figure supplement 1C). In order to make sure that the scramblase is surrounded by a sufficiently large membrane area, we assembled 2N2 scaffolding proteins with nhTMEM16 at three different lipid to protein ratios (LPR) in the absence of Ca^2+^ and collected small datasets for all of them (Figure 2-figure supplement 1). In all cases, the protein is located close to the center of an elliptic disc with its long dimension oriented about parallel to the shortest side of the disc, a behavior that is more pronounced in larger discs obtained at high lipid to protein ratios (Figure 2-figure supplement 1). The orientation reflects a preference of lipids for locations distant from the subunit cavity, which appears to deform the nanodisc. Since increasing the amount of lipids only caused elongation of the nanodisc in one direction without a change in its diameter near the subunit cavity, we proceeded with data collection on the sample with the lowest LPR of 145:1, which resulted in a homogeneous assembly of approximately 165 x 140 Å in size (Figure 2-figure supplement 1A). This approach was used to determine the structures of ligand-bound and ligand-free states of nhTMEM16 in a membrane-like environment. The data is of excellent quality and provide an unambiguous view of nhTMEM16 in different conformations (Figures 2 and 3; Figure 2-figure supplements 2; Figure 3-figure supplements 1-2; Tables 1 and 2).

**Table 2.** Cryo-EM data collection, refinement and validation statistics of the Ca^2+^ bound nhTMEM16 in nanodiscs.

|  | nhTMEM16,<br>2N2, +Ca<sup>2+</sup><br>open<br>(EMDB-4592,<br>PDB 6QM9) | nhTMEM16,<br>2N2, +Ca<sup>2+</sup><br>intermediate-closed<br>(EMDB-4593,<br>PDB 6QMA) | nhTMEM16,<br>2N2, +Ca<sup>2+</sup><br>closed<br>(EMDB-4594,<br>PDB 6QMB) |
| --- | --- | --- | --- |
| **Data collection and processing** |  |  |  |
| Microscope |  | FEI Talos Arctica |  |
| Camera |  | Gatan K2 Summit + GIF |  |
| Magnification |  | 49,407 |  |
| Voltage (kV) |  | 200 |  |
| Exposure time frame/total (s) |  | 0.15/9 |  |
| Number of frames per image |  | 60 |  |
| Electron exposure (e-/Å<sup>2</sup>) |  | 52 |  |
| Defocus range (µm) |  | -0.5 to -2.0 |  |
| Pixel size (Å) |  | 1.012 |  |
| Box size (pixels) |  | 240 |  |
| Symmetry imposed |  | C2 |  |
| Initial particle images (no.) |  | 2,440,110 |  |
| Final particle images (no.) | 71,175 | 33,310 | 41,631 |
| Map resolution (Å) |  |  |  |
| 0.143 FSC threshold | 3.57 | 3.68 | 3.57 |
| Map resolution range (Å) | 3.4 – 5 | 3.6 – 5 | 3.4 – 5 |
| **Refinement** |  |  |  |
| Initial model used | 6QM5 | 6QMB | 6QM5 |
| Model resolution (Å) | 3.6 | 3.7 | 3.6 |
| FSC threshold |  |  |  |
| Model resolution range (Å) |  |  |  |
| Map sharpening *B* factor (Å<sup>2</sup>) | -114 | -96 | -95 |
| Model composition |  |  |  |
| Nonhydrogen atoms | 10878 | 10714 | 10696 |
| Protein residues | 1346 | 1324 | 1322 |
| Ligands | 4 | 4 | 4 |
| *B* factors (Å<sup>2</sup>) |  |  |  |
| Protein | 77.93 | 84.86 | 88.53 |
| Ligand | 54.34 | 57.57 | 67.10 |
| R.m.s. deviations |  |  |  |
| Bond lengths (Å) | 0.005 | 0.006 | 0.008 |
| Bond angles (°) | 0.880 | 0.872 | 0.858 |
| Validation |  |  |  |
| MolProbity score | 1.36 | 1.38 | 1.42 |
| Clashscore | 3.74 | 4.69 | 3.90 |
| Poor rotamers (%) | 0 | 0.18 | 0 |
| Ramachandran plot |  |  |  |
| Favored (%) | 96.82 | 97.23 | 96.38 |
| Allowed (%) | 3.18 | 2.77 | 3.62 |
| Disallowed (%) | 0 | 0 | 0 |

The differences between structures of nhTMEM16 determined in detergents and in a lipid environment are most pronounced for the Ca^2+^-free state determined at 3.8 Å resolution (Figure 2A,B,D; Figure 2-figure supplement 2). Whereas the structure in detergent displays a Ca^2+^-free conformation where the subunit cavity remains open to the membrane, the same region has changed its accessibility in the nanodisc sample. Here, the movement of α4 leads to a shielding of the cavity from the membrane core, resembling a conformation observed in the ion channel TMEM16A (Figure 2E) (Dang et al., 2017; Paulino et al., 2017a; Paulino et al., 2017b). This movement is accompanied by a conformational change of α6 as a consequence of the repulsion of negatively charged residues in the empty Ca^2+^-binding site (Figure 2B-D). Generally, the Ca^2+^-free state of nhTMEM16 in nanodiscs is very similar to equivalent structures of its close homologs afTMEM16 (Falzone et al., 2019), TMEM16K (Bushell et al., 2018) and the more distantly related scramblase TMEM16F (Alvadia et al., 2018) (Figure 2C,F and Figure 1-figure supplement 4), thus underlining that all structures represent inactive conformations of the scramblase in the absence of ligand.

**Figure 2.**
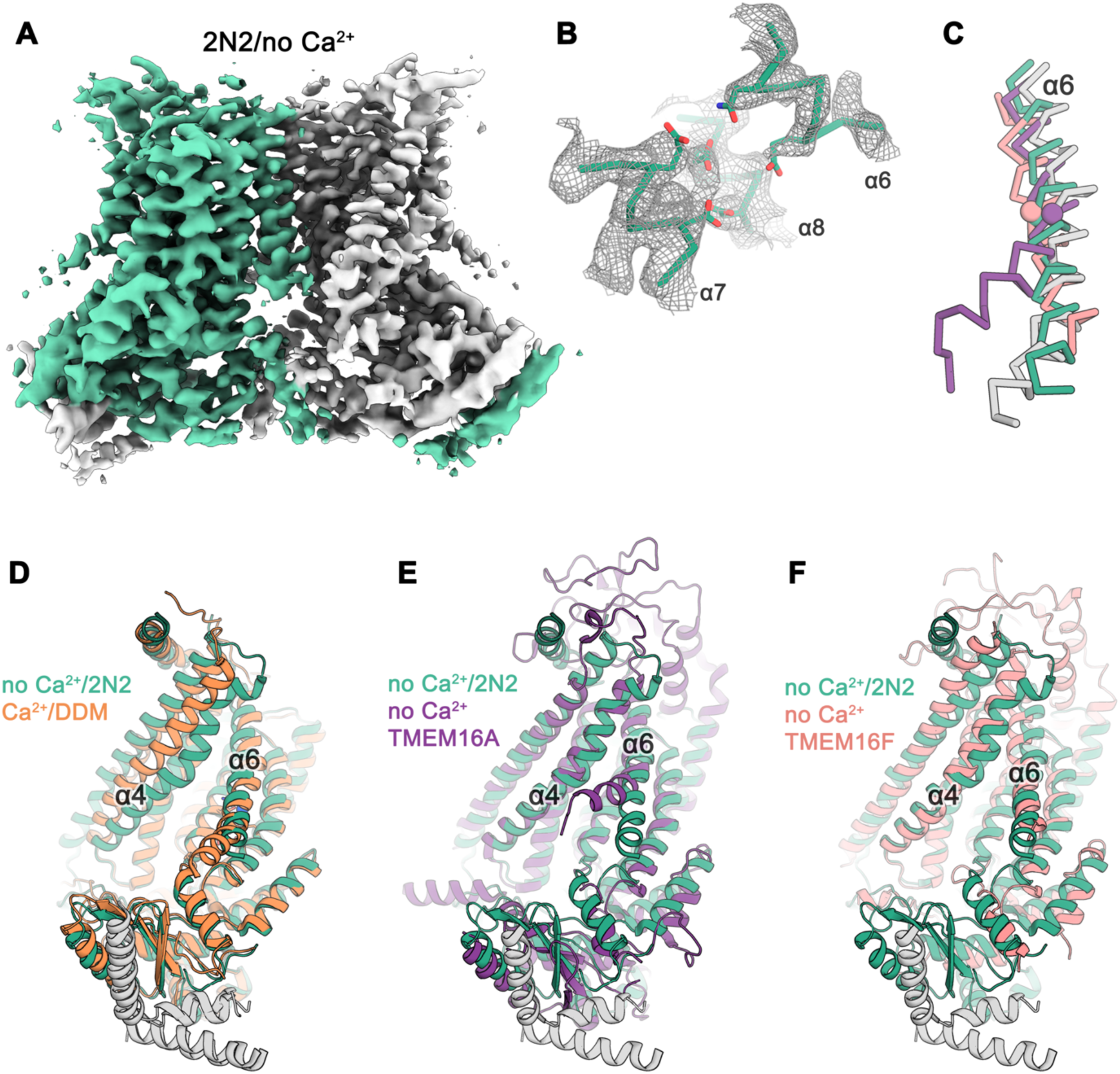
Cryo-EM structure of nhTMEM16 in nanodiscs in absence of Ca^2+^. (**A**) Cryo-EM map of the nhTMEM16 dimer (light green and grey) in nanodiscs in absence of Ca^2+^ at 3.8 Å resolution. The map was sharpened with a b-factor of –150 Å^2^ and is contoured at 5.6 σ. (**B**) View of the Ca^2+^-binding site in the Ca^2+^-free state of nhTMEM16 in nanodiscs. The cryo-EM density shown as mesh is contoured at 6 σ, the backbone is displayed as Cα-trace and selected side-chains as sticks. (**C**) Cα-traces of α6 from a superposition of Ca^2+^-free structures of nhTMEM16 in nanodiscs (green), TMEM16A (violet, PDBID: 5OYG) and TMEM16F (red) and of the Ca^2+^-bound structure of nhTMEM16 in DDM (grey). The spheres indicate the position of the flexible glycine residue in TMEM16A and TMEM16F, which acts as a hinge for conformational changes. (**D-F**) Ribbon representation of superpositions of the Ca^2+^-free structure in nanodiscs (light green and grey) and the Ca^2+^-bound structure in DDM (**D**, orange and grey), the Ca^2+^-free structure of TMEM16A (**E**, violet, PDBID: 5OYG), the Ca^2+^-free structure of TMEM16F (**F**, red). Selected α-helices are labeled, the views are as in Figure 1.

### Structures of Ca^2+^-bound states of nhTMEM16 in lipid nanodiscs

Different from the data of the Ca^2+^-free protein in nanodiscs, which comprises a single prevailing state, the cryo-EM data obtained for the Ca^2+^-bound samples in the same environment shows a large heterogeneity along the ‘subunit cavity’. This heterogeneity allows to sample the transition of the lipid permeation path from the closed to the open state and emphasizes the intrinsic dynamics of this region during activation (Figure 3; Figure 3-figure supplements 1 and 2; Table 2). A classification of the data yielded several states with distinct conformations of the ‘subunit cavity’, all of which exhibit well-resolved cryo-EM density for two Ca^2+^ ions and a rigid arrangement of α6 in place to coordinate the divalent ions (Figure 3A-C; Figure 3-figure supplements 1 and 2). About 50% of the particles define classes where α3 and α4 are poorly resolved, thus emphasizing the high degree of flexibility in the region, which suggest that several states along the activation process might be transient (classes 5 and 6, Figure 3-figure supplement 1C). By contrast, three classes show well resolved distinct conformations. The ‘open’ class, at 3.6 Å resolution, encompasses about 25% of the particles and strongly resembles the Ca^2+^-bound open state obtained in detergent, with a subunit cavity that is exposed to the membrane (Figure 3A; Figure 3-figure supplements 1 and 2). In contrast, the ‘Ca^2+^-bound closed’ class determined from 15% of particles at 3.6 Å resolution, strongly resembles the ‘Ca^2+^-free closed’ state obtained in nanodiscs, with a subunit cavity that is shielded from the membrane, except that Ca^2+^ is bound and α-helix 6 is less mobile (Figure 3C,G; Figure 3-figure supplements 1 and 2). The third distinct state at 3.7 Å resolution encompasses 12% of all particles and shows an ‘intermediate’ conformation, where the subunit cavity is still shielded from the membrane but where a potential aqueous pore surrounded by α-helices of the subunit cavity has widened compared to the closed state (Figure 3B,E,F; Figure 3-figure supplements 1 and 2). The distribution of states, with a portion of nhTMEM16 residing in potentially non-conductive conformations correlates with the lower activity of the protein in this lipid composition (Figure 1-figure supplement 1C) and it emphasizes the equilibrium between conformations of nhTMEM6 in the Ca^2+^-bound state. Unassigned weak density in the subunit cavity in the ‘open’ - but not in the ‘closed’ - conformation, hints at lipid molecules during the transition between the leaflets (Figure 3-figure supplement 3). Strikingly, the strongest density is located at an equivalent position as found for the potential DDM density in the cryo-EM maps of nhTMEM6 in detergent (Figure 1-figure supplement 5).

**Figure 3.**
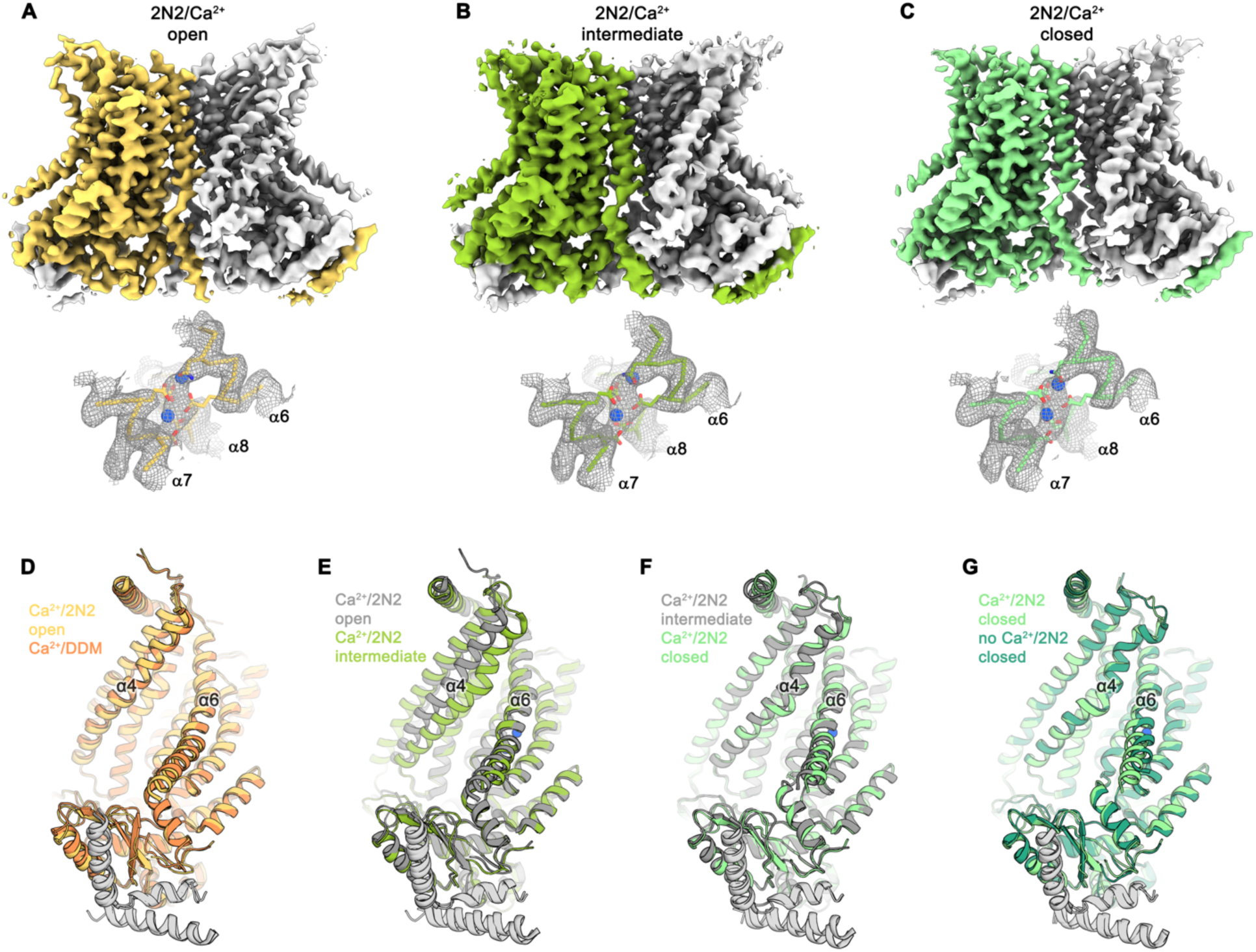
Cryo-EM structures of nhTMEM16 in nanodiscs in presence of Ca^2+^. (**A**) Cryo-EM map of the nhTMEM16 dimer in the Ca^2+^-bound ‘open’ state (yellow and grey) in nanodiscs at 3.6 Å sharpened with a b-factor of –114 Å^2^ and contoured at 6.5 σ (upper panel). Close-up of the Ca^2+^-binding site with two bound Ca^2+^-ions displayed as blue spheres (lower panel). Cryo-EM density contoured at 7 σ is shown as grey mesh, the backbone is displayed as yellow Cα-trace, selected side-chains as sticks. (**B**) Cryo-EM map of the nhTMEM16 dimer in the Ca^2+^-bound ‘intermediate’ state (yellow-green and grey) in nanodiscs at 3.7 Å sharpened with a b-factor of – 96 Å^2^ and contoured at 4.9 σ (upper panel). Close-up of the Ca^2+^-binding site with two bound Ca^2+^-ions displayed as blue spheres (lower panel). Cryo-EM density contoured at 7 σ is shown as grey mesh, the backbone is displayed as light green Cα-trace, selected side-chains as sticks. (**C**) Cryo-EM map of the nhTMEM16 dimer in the ‘Ca^2+^-bound closed state (green and grey) in nanodiscs at 3.6 Å sharpened with a b-factor of –96 Å^2^ and contoured at 6.5 σ (upper panel). Close-up of the Ca^2+^-binding site with two bound Ca^2+^-ions displayed as blue spheres (lower panel). Cryo-EM density contoured at 7 σ is shown as grey mesh, the backbone is displayed as light green Cα-trace, selected side-chains as sticks. (**D-G**) Ribbon representation of superpositions with Ca^2+^-bound structures of nhTMEM16 in nanodiscs: (**D**) Ca^2+^-bound open structure in nanodiscs and Ca^2+^-bound structure in DDM; (**E**) Ca^2+^-bound open structure in nanodiscs and Ca^2+^-bound intermediate structure in nanodiscs; (**F**) Ca^2+^-bound intermediate structure in nanodiscs and Ca^2+^-bound closed structure in nanodiscs; (**G**) Ca^2+^-bound closed structure in nanodiscs and Ca^2+^-free structure in nanodiscs. Selected helices are labeled and views are as in Figure 1.

### Conformational transitions

The diversity of conformations of nhTMEM16 observed in various datasets and the presence of different populations in the Ca^2+^-bound sample in lipid nanodiscs suggest that our structures represent distinct states along a stepwise activation process. Apart from a small rearrangements of the cytosolic domain, the core of the protein is virtually identical in all structures and the largest movements are observed in the vicinity of the subunit cavity (Figures 2D, 3D-G 4 and 5A and Video 1). The Ca^2+^-bound conformation in detergent and the ‘open’ class observed in the data of Ca^2+^-bound protein in nanodiscs, both show the widest opening of the subunit cavity and thus likely depict a scrambling-competent state of the protein (Figure 4A). In this open state, α-helices 4 and 6 are separated from each other on their entire length framing the two opposite edges of a semicircular polar furrow that is exposed to the lipid bilayer (Figure 4A). In this case, the interaction of α3 with α10 of the adjacent subunit on the intracellular side appears to limit the movement of α4, thus preventing a further widening of the cavity on the intracellular side (Figures 5A-C). On the opposite border of the subunit cavity, α6 is immobilized in its position by the bound Ca^2+^ ions, which results in a tight interaction with α7 and α8 (Figure 5D). In the ‘intermediate’ class found in the Ca^2+^-bound sample in nanodiscs, we find marked conformational changes compared to the ‘open state’. The most pronounced differences are found for the α3-α4 pair. The intracellular part of α3 (but not the α2-α3 loop contacting the helix Cα1 following α10) has detached from its interaction with α10, leading to a concomitant movement of α4 at the same region (Figure 5B). This movement has also affected the conformation of the extracellular part of α4, which has rotated and approached α6, thereby forming initial contacts between both helices (Figures 4B and 5C,D). The subunit cavity in this conformation is no longer exposed to the membrane at its extracellular part and the structure likely represents an intermediate of the scramblase towards activation (Figure 4B). This ‘intermediate state’ is remarkable in light of the observed ion conduction of nhTMEM16 and other TMEM16 scramblases (Alvadia et al., 2018; Lee et al., 2016; Suzuki et al., 2013; Yang et al., 2012; Yu et al., 2015), as it might harbor a potential protein-enclosed aqueous pore located in region that is expanded compared to the ‘closed state’ (Figure 5-figure supplement 1). The interactions between α4 and α6 on the extracellular side further tighten in the ‘closed’ class found in the Ca^2+^-bound sample in nanodiscs, thereby constricting the aqueous pore observed in the ‘intermediate state’ (Figures 4C; Figure 5-figure supplement 1). This conformation strongly resembles the Ca^2+^-free ‘closed’ conformation observed in nanodiscs, except that in the latter, the dissociation of the ligand from the protein has allowed a relaxation of α6 leading to its detachment from α8 at the intracellular part of the helix (Figures 4D and 5C,D). Although, unlike TMEM16A and TMEM16F, nhTMEM16 does not contain a flexible glycine residue at the hinge, the movements of α6 upon Ca^2+^-release occur at a similar region (Alvadia et al., 2018; Paulino et al., 2017a) (Figure 2C-F and Figure 1-figure supplement 4). In the transition from an ‘open’ to a ‘closed state’, α4 undergoes the largest changes among all parts of nhTMEM16 with two prolines (Pro 332 and Pro 341) and a glycine (Gly 339) serving as potential pivots for helix-rearrangements (Figure 5E and Video 1). Remarkably, both prolines are conserved among fungal homologues and Pro 332 is also found in TMEM16K (Bushell et al., 2018), the closest mammalian orthologue of nhTMEM16 (Figure 1-figure supplement 4).

**Figure 4.**
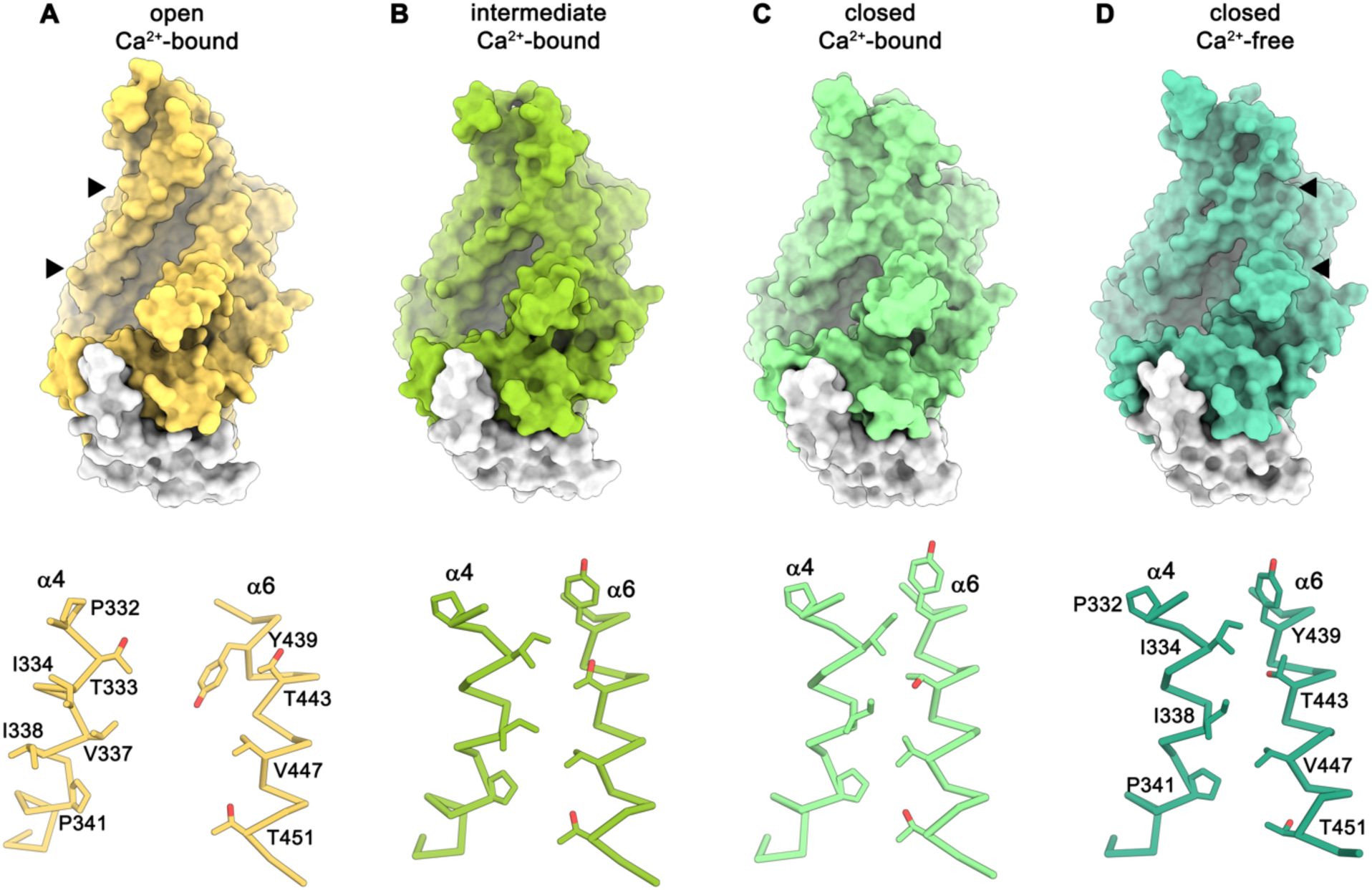
Conformations of the ‘subunit cavity’. Views of the molecular surface of the subunit cavity (top) and structures of α4 and α6 that line the cavity displayed as Cα-trace with selected sidechains shown as sticks and labeled (bottom). (**A**) ‘Open state’ as defined in the Ca^2+^-bound structure in nanodiscs. (**B**) ‘Intermediate state’ as defined in the Ca^2+^-bound structure in nanodiscs, (**C**) ‘Ca^2+^-bound closed state’ as defined in the Ca^2+^-bound structure in nanodiscs. (**D**) ‘Closed state’ as defined by the Ca^2+^-free structure in nanodiscs. The region shown in the lower panels is depicted by triangles in the upper panels.

**Figure 5.**
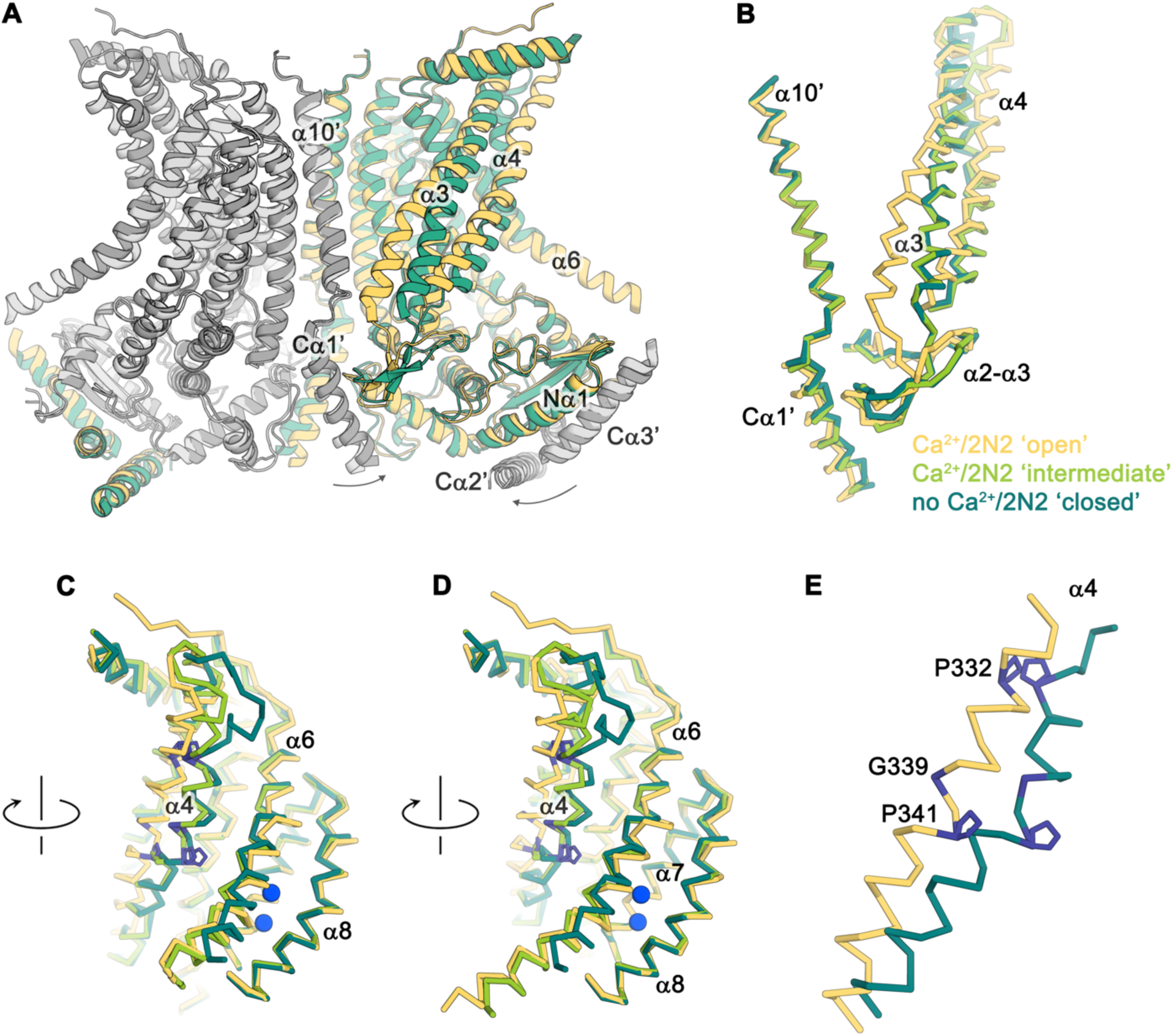
Conformational changes. (**A**) Ribbon representation of a superposition of the Ca^2+^-bound ‘open’ (yellow and light grey) and Ca^2+^-free ‘closed’ (green and dark grey) nhTMEM16 structures in nanodiscs. Selective helices are labeled and the arrows indicate small rearrangements of the cytosolic domain upon Ca^2+^-release. (**B**-**D**) Cα-traces of selected regions of the ‘subunit cavity’ in different conformations as defined by the Ca^2+^-bound ‘open’ structure in nanodiscs (yellow), the Ca^2+^-bound ‘intermediate’ structure in nanodiscs (light green) and the Ca^2+^-free ‘closed’ structure in nanodiscs (dark green). The view in B is as in A and the respective orientations of subsequent panels are indicated. (**D**) Superposition of α4 in the ‘open’ and closed’ states with residues that act as potential pivots for structural arrangements highlighted in blue.

### Protein-induced distortion of the lipid environment

A remarkable feature observed in the cryo-EM maps concerns the arrangement of detergent or lipid molecules surrounding nhTMEM16, which allowed us to characterize the influence of the protein on its environment. The observed distortions are found to a similar extent in all determined structures, irrespective of the presence or absence of calcium in detergent or lipid nanodiscs and can thus be considered a state-independent property of the protein. Neither detergent molecules nor nanodisc lipids are uniformly distributed on the outside of nhTMEM16 (Figure 6 and Video 2). Instead, they adapt to the shape of the protein with distortions observed in the vicinity of the shorter α-helices 1 and 8 at the extracellular side and at the gap between α-helices 4 and 6 at the intracellular side (Figure 6). These distortions result in a marked deviation from the annular shape of detergents molecules or lipids surrounding the protein found in most membrane proteins. Remarkably, the resulting undulated distribution contains depressions in both the detergent and lipid density close to the entrances of the subunit cavity (Figure 6 and Video 2). The arrangement of the protein with its long dimension parallel to the short diameter of the oval-shaped nanodiscs reflects the preferential location of lipids at sites distant from the subunit cavity (Figure 2-figure supplement 1). The protein affects the distribution of lipids and the shape of the plane of the bilayer in nanodiscs, which is slightly V-shaped and locally deviates from planarity at several regions (Figure 6B,C). These pronounced depressions thin and distort the lipid structure at both ends of the subunit cavity, thereby likely facilitating the entry of polar head-groups into the cavity and lowering the barrier for lipid movement (Figure 6B).

**Figure 6.**
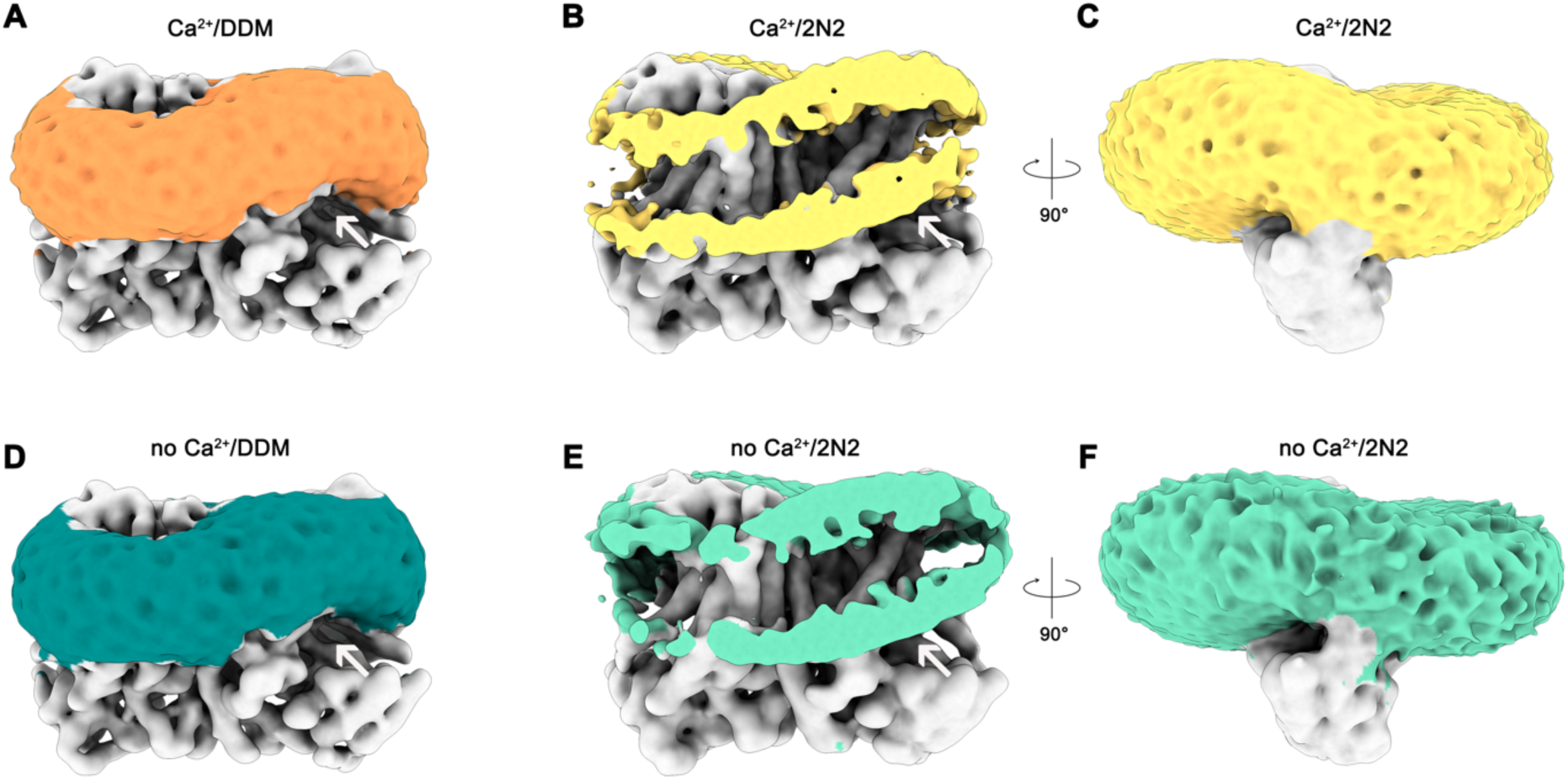
Detergent and lipid interactions. Refined and unmasked cryo-EM density map low-pass filtered to 6 Å are shown. (**A**) Map of the Ca^2+^-bound nhTMEM16 data in detergent contoured at 4 σ; (**B**,**C**) Map of the Ca^2+^-bound nhTMEM16 data in nanodiscs contoured at 3.2 and 1.6 σ; Map of the Ca^2+^-free nhTMEM16 data in detergent contoured at 4 σ; (**E** and **F**), Map of the Ca^2+^-free nhTMEM16 data in nanodiscs contoured at 3.2 and 1.6 σ. The density corresponding to the detergent micelle or the nanodisc, which is composed of lipids surrounded by the 2N2 belt protein, are colored in orange, yellow, dark green and light green, respectively, density of nhTMEM16 is shown in grey. A,B,D,E show a front view of the dimer similar to Figure 1A and the subunit cavity is indicated by a white arrow. B, E, clipped maps reveal the headgroup regions of both membrane leaflets. C,F, are viewed in direction of the subunit cavity.

### Functional properties of mutants affecting the activation process

The distinct conformations of nhTMEM16 determined in this study hint at a sequential mechanism, in which Ca^2+^-dependent and independent steps are coupled to promote activation similar to ligand-dependent ion channels. We have thus investigated the functional consequences of mutations of residues of the Ca^2+^-binding site and of the hinges for α4 movement with our liposome-based lipid scrambling assay. The mutation D503A located on α7 concerns a residue of the Ca^2+^-binding site that does not change its position in different protein conformations and thus should affect the initial ligand-binding step (Figure 7A). In this case, we find a Ca^2+^-dependent scrambling activity, although with strongly decreased potency of the ligand (Figures 7B,C). Next, we investigated whether the mutation of two proline residues and a close-by glycine (Pro 332, Gly 339 and Pro 341), which form potential pivots during the rearrangement of α4 would affect lipid scrambling (Figure 7A). Whereas we did not find any detectable effect in the mutant P341A, the mutants G339A and P332A showed a strongly decreased activity both in the presence and absence of Ca^2+^ (but no detectable change in the Ca^2+^-potency of the protein, Figures 7D-F; Figure 7-figure supplement 1). As the residues do not face the subunit cavity in the ‘open state’ and are unlikely to interact with translocating lipids, we assume that these mutants alter the equilibrium between active and inactive conformations thus emphasizing the role of conformational changes in α-helix 4 for nhTMEM16 activation.

**Figure 7.**
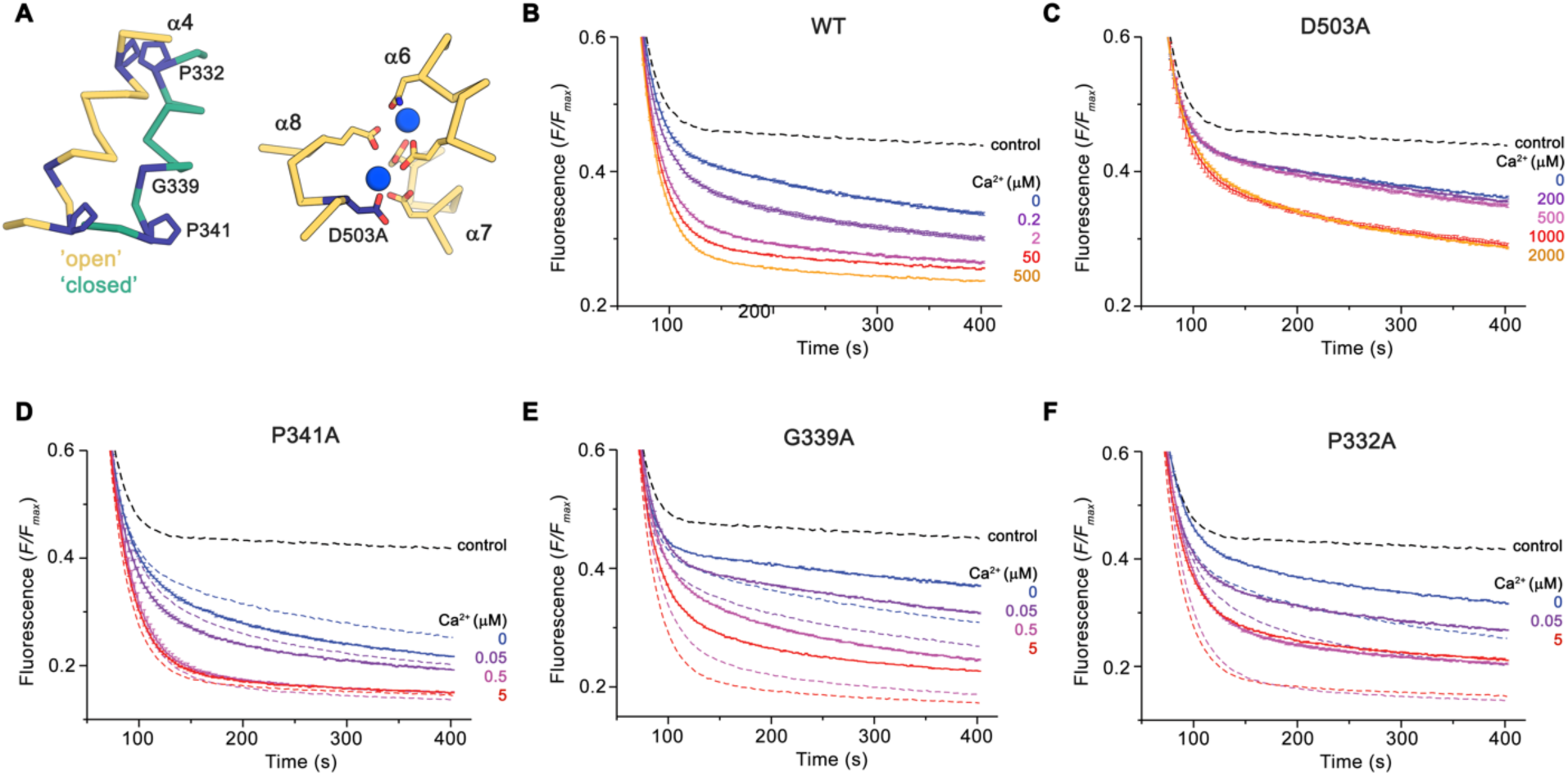
Functional properties of mutants. (**A**) Regions of nhTMEM16 harboring mutated residues. Left, Cα-trace of α4 in ‘open’ (yellow) and ‘closed’ (green) conformations of Ca^2+^-bound nhTMEM16 in nanodiscs. Right, ion binding site with Ca^2+^-ions shown in blue. (**B**-**F**) Ca^2+^-dependence of scrambling activity in nhTMEM16-containing proteoliposomes. Traces depict sections of the fluorescence decrease of tail-labeled NBD-PE lipids after addition of dithionite at different Ca^2+^ concentrations. Data show averages of three technical replicates, errors are s.e.m.. Ca^2+^ concentrations (µM) are indicated. Full traces are displayed in Fig7-figure supplement 1. (B) WT, (C) D503A, (D) P341A, (E) G339A, and (F) P332A. (D to F) The traces of WT reconstituted in the same batch of liposomes are shown as dashed lines in the same color as their corresponding conditions of the mutant for comparison. (B to F), The black dashed line refers to the fluorescence decay in protein-free liposomes.

## Discussion

By combining data from cryo-electron microscopy and biochemical assays, our study has addressed two important open questions concerning the function of a TMEM16 scramblase: it has revealed the conformational changes that lead to the activation of the protein in response to Ca^2+^ binding and it has shown how the protein interacts with the surrounding membrane to facilitate lipid translocation. Structures in Ca^2+^-free and bound states reveal distinct conformations of nhTMEM16, which capture the catalytic subunit cavity of the scramblase at different levels of exposure to the membrane during the activation process, representing a ‘closed’, at least one ‘intermediate’ and an ‘open state’. As different protein conformations are observed in presence and absence of its ligand, we propose a stepwise activation mechanism for TMEM16 scramblases, where all states are at equilibrium (Figure 8; Figure 8-figure supplement 1). Here, the Ca^2+^-free conformation obtained in nanodiscs defines a non-conductive state of the protein, where the polar subunit cavity is shielded from the membrane by tight interactions of α4 and α6 on the extracellular part of the membrane (Figure 4D). The protein surface of the closed subunit cavity is hydrophobic as expected for a membrane protein. This ‘closed state’ of nhTMEM16 resembles equivalent structures obtained for the closely related lipid scramblases afTMEM16 (Falzone et al., 2019), TMEM16K (Bushell et al., 2018), the more distantly related TMEM16F (Alvadia et al., 2018) and the ion channel TMEM16A (Figure 2E,F) (Dang et al., 2017; Paulino et al., 2017a; Paulino et al., 2017b), thus underlining that it is representative for an inactive conformation of both functional branches of the family. In this conformation, the Ca^2+^-binding site is empty, with the intracellular part of α6 positioned apart from the remainder of the binding site to release the electrostatic strain induced by the large negative net-charge in this region. Details of the conformation of α6 in the absence of Ca^2+^ differ between the family members of known structure with the conformation in nhTMEM16 being closer to the one observed for TMEM16F (Alvadia et al., 2018; Paulino et al., 2017a).

**Figure 8.**
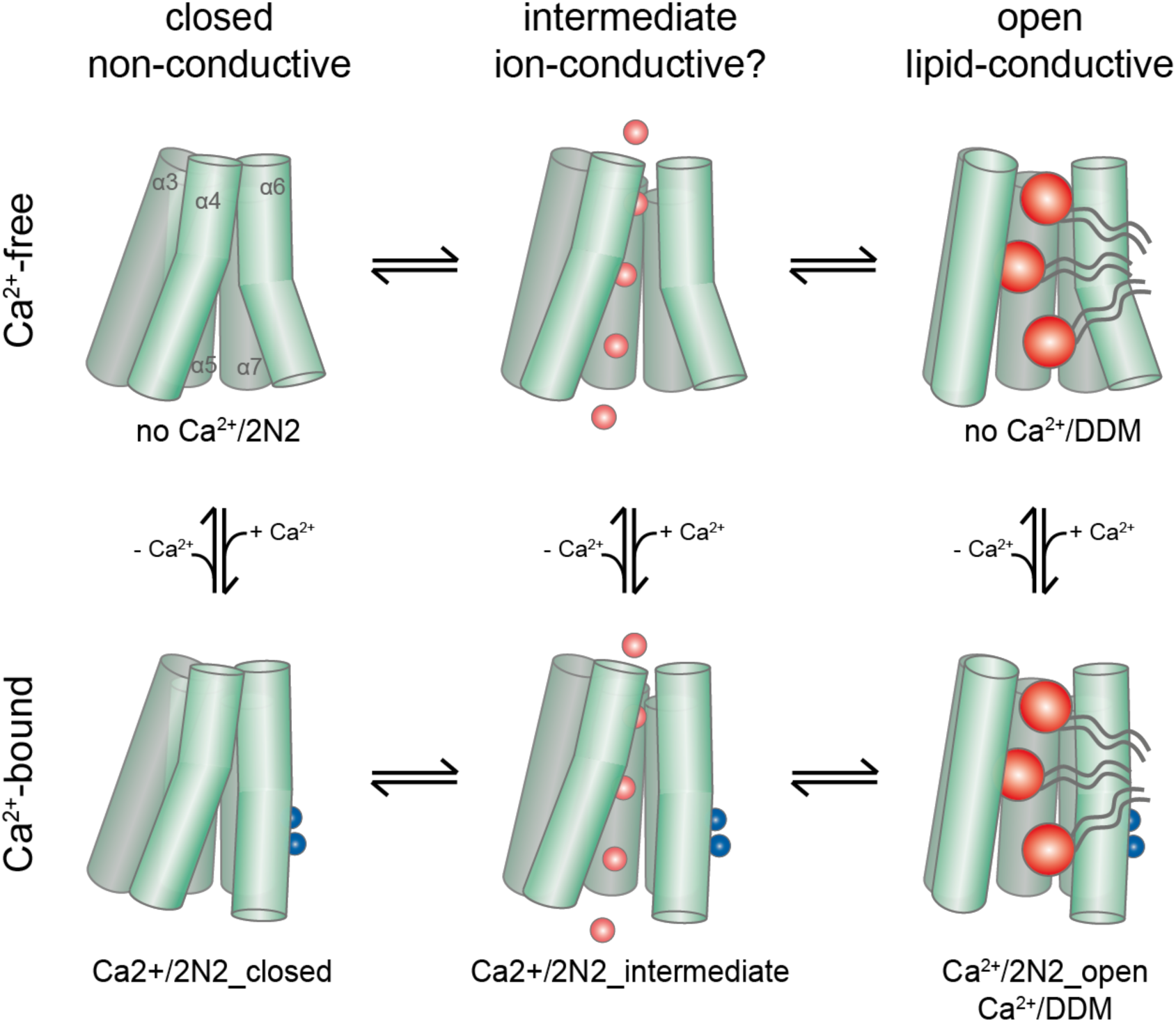
Activation mechanism. Scheme of the stepwise activation of nhTMEM16 displaying the equilibrium of states in Ca^2+^-bound and Ca^2+^-free conditions. Conformations obtained in this study and their correspondence to distinct states are indicated. Ca^2+^ and permeating ions are depicted as blue and red spheres, respectively. Phospholipid headgroups are shown as red spheres and acyl chains as grey tails.

During the activation process, Ca^2+^ binding provides interactions with residues on α6. The helix rearranges into a locked state, as manifested in the well resolved cryo-EM density for its intracellular half, which shields the binding site from the cytoplasm. This Ca^2+^-induced movement of α6 in nhTMEM16 is coupled to rearrangements in its interface with α4, potentially weakening the interactions and resulting in a widening of the cavity. A cavity with a changed accessibility to the membrane is displayed in a population of particles observed in the Ca^2+^-bound nhTMEM16 structure in nanodiscs (Figure 4B). As the subunit cavity is not yet exposed to the membrane, the ‘intermediate state’ likely does not promote lipid scrambling. Instead, it might have opened a protein-enclosed pore that could facilitate ion conduction (Figure 5-figure supplement 1), which has been described as by-product of lipid scrambling in nhTMEM16 (Lee et al., 2016) and which is a hallmark of the lipid scramblase TMEM16F (Alvadia et al., 2018; Yang et al., 2012; Yu et al., 2015). Following the initial Ca^2+^-induced conformational transition, the activation of lipid scrambling requires a second step, which leads to a larger reorientation of α-helices 3 and 4 and a subsequent opening of the polar subunit cavity to the membrane (Figures 3 and 4). This open state is defined by the Ca^2+^-bound structure of nhTMEM16 obtained in detergent and by one of the classes observed in the Ca^2+^-bound nanodisc dataset.

Thus, ion conduction and lipid-scrambling in nhTMEM16 and other TMEM16 scramblases might be mediated by distinct conformations which are at equilibrium in a Ca^2+^-bound protein (Figure 8; Figure 8-figure supplement 1). This stepwise activation mechanism is in general accordance with molecular dynamics simulations, which have started from the fully Ca^2+^-bound open structure and for which transitions leading to a partial closure of the subunit cavity have been observed in the trajectories (Jiang et al., 2017; Lee et al., 2018). The basal scrambling activity of nhTMEM16 is likely a consequence of the equilibrium between open, intermediate and closed conformations, which is strongly shifted towards the closed state in the absence of Ca^2+^. This is consistent with the observation of a Ca^2+^-free open state in detergent but not in nanodiscs, where the detergent environment favors the equilibrium towards the open conformation (Figure 1H).

The cryo-EM structures of nhTMEM16 also reveal how the protein interacts with the surrounding membrane to disturb its structure and consequently lower the energy barrier for lipid flip-flop, as predicted from molecular dynamics simulations (Bethel and Grabe, 2016; Jiang et al., 2017; Lee et al., 2018; Stansfeld et al., 2015). Our data reveal a pronounced distortion of the protein-surrounding environment, which is stronger than observed for the ion channel TMEM16A (Paulino et al., 2017b). This effect is preserved in all four datasets, irrespectively of the presence or absence of ligand in detergent or nanodiscs, as their underlying structural features change little in the different conformations. The distortion causes a deviation of the detergent or lipid molecules surrounding nhTMEM16 from the annular structure observed in most membrane proteins (Figure 6 and Video 2). The resulting deformation of the membrane at the entrances to the subunit cavity potentially serves to channel lipid headgroups through this hydrophilic furrow on their way across the membrane.

In summary, our structures of nhTMEM16 have described how a calcium-activated lipid scramblase is activated in a stepwise manner and how it bends the surrounding bilayer to promote the transport of lipids between both leaflets of the membrane by a mechanism that is likely conserved within the family.

## Materials and methods

### Key resources table

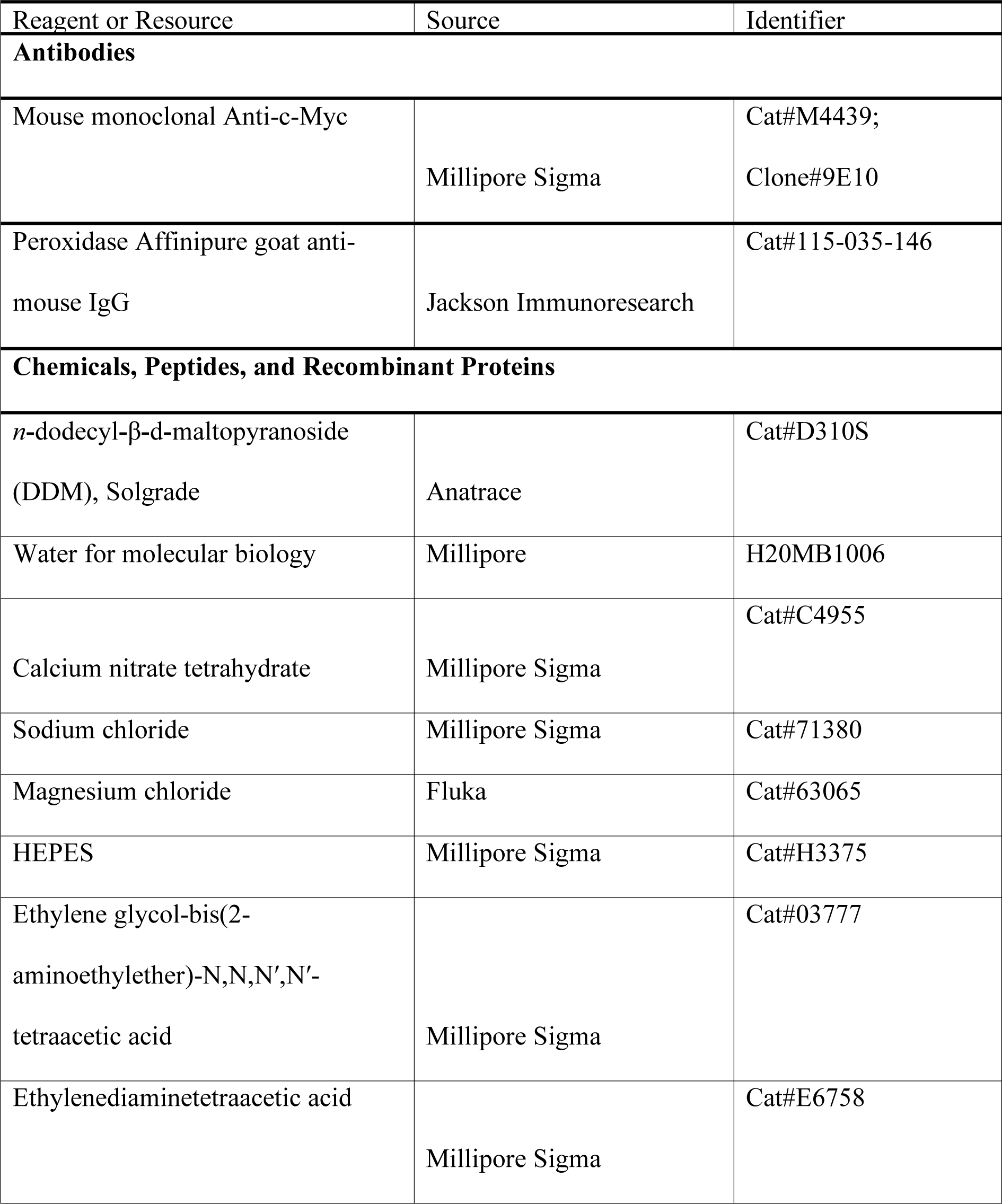

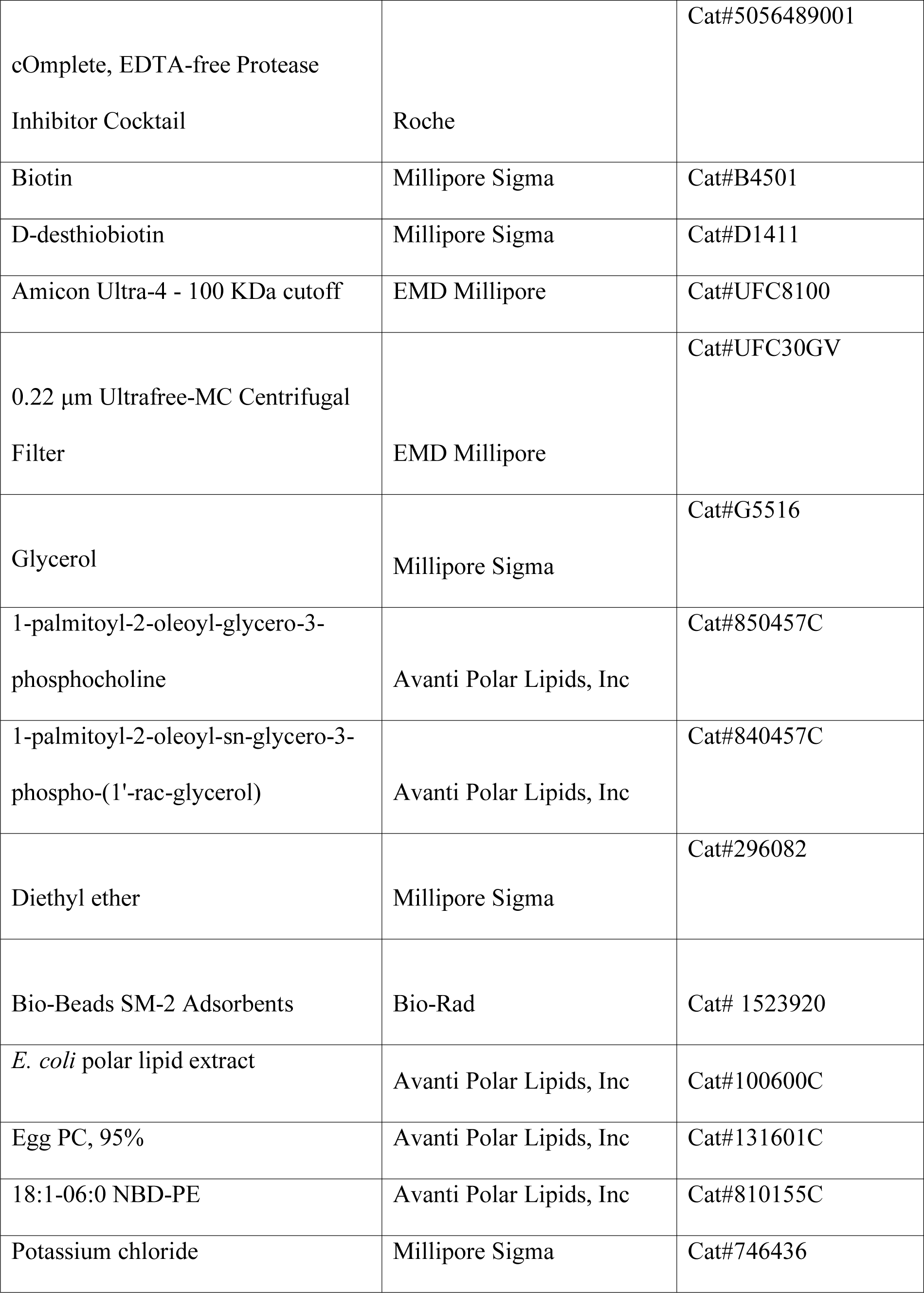

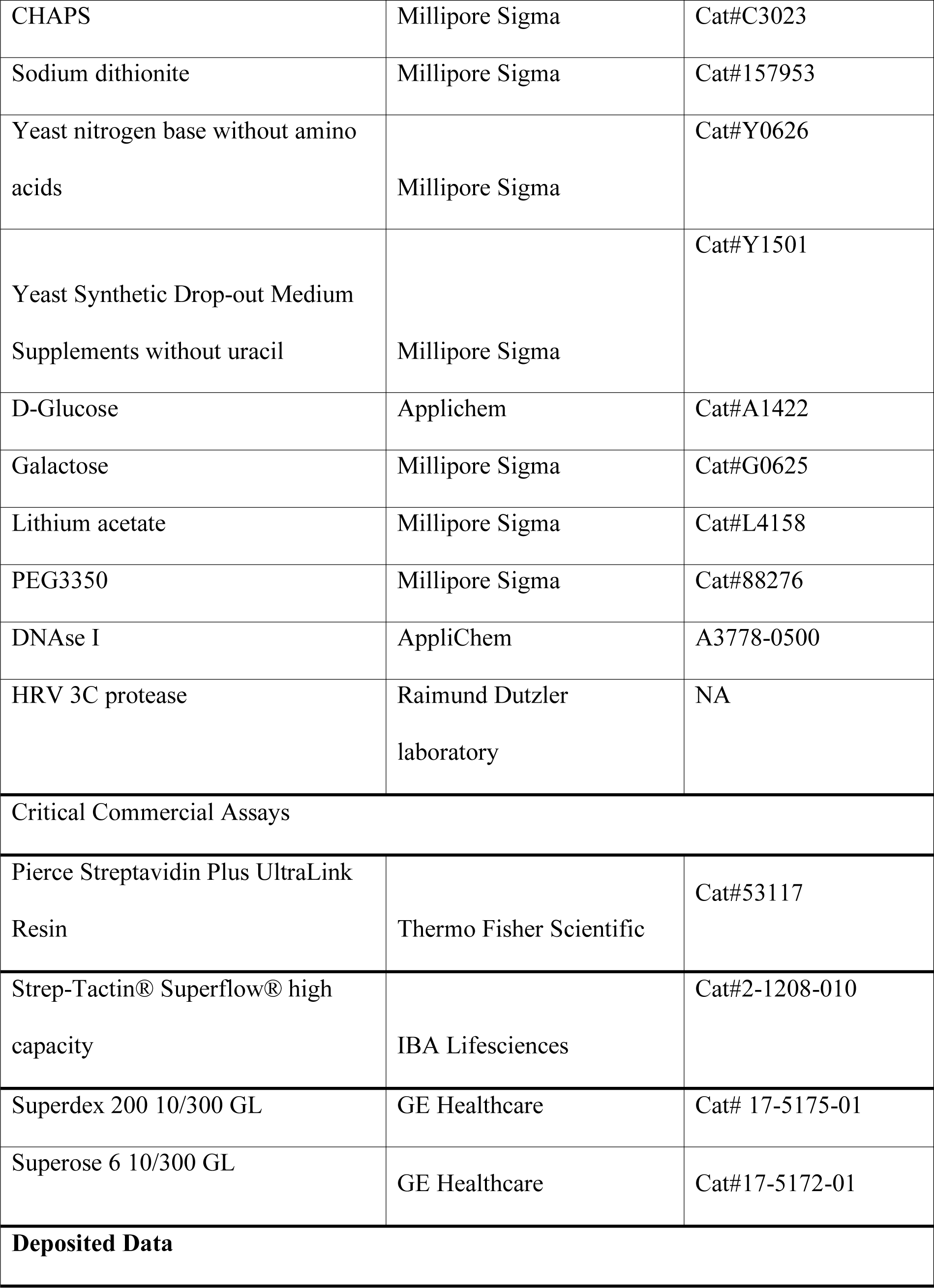

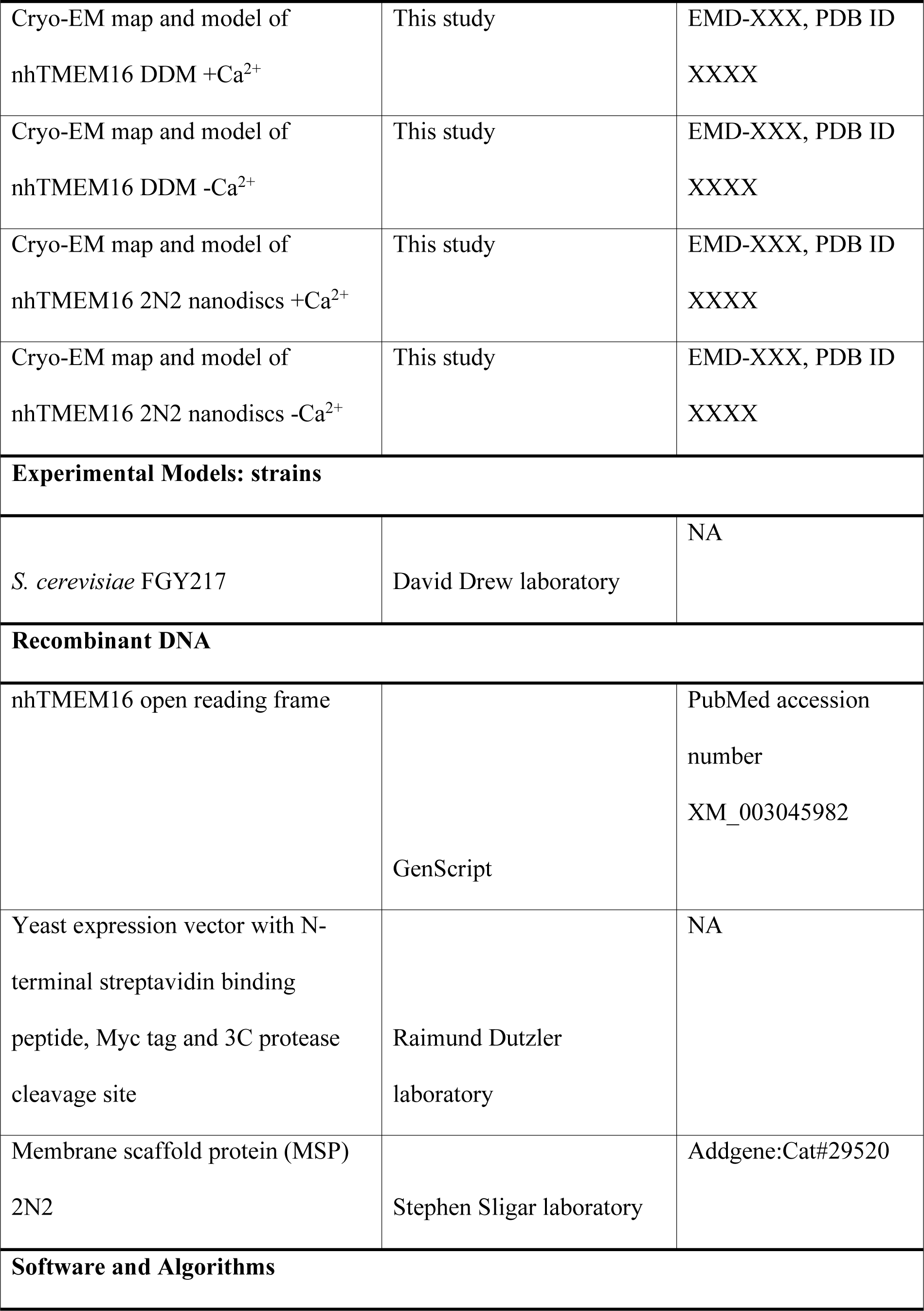

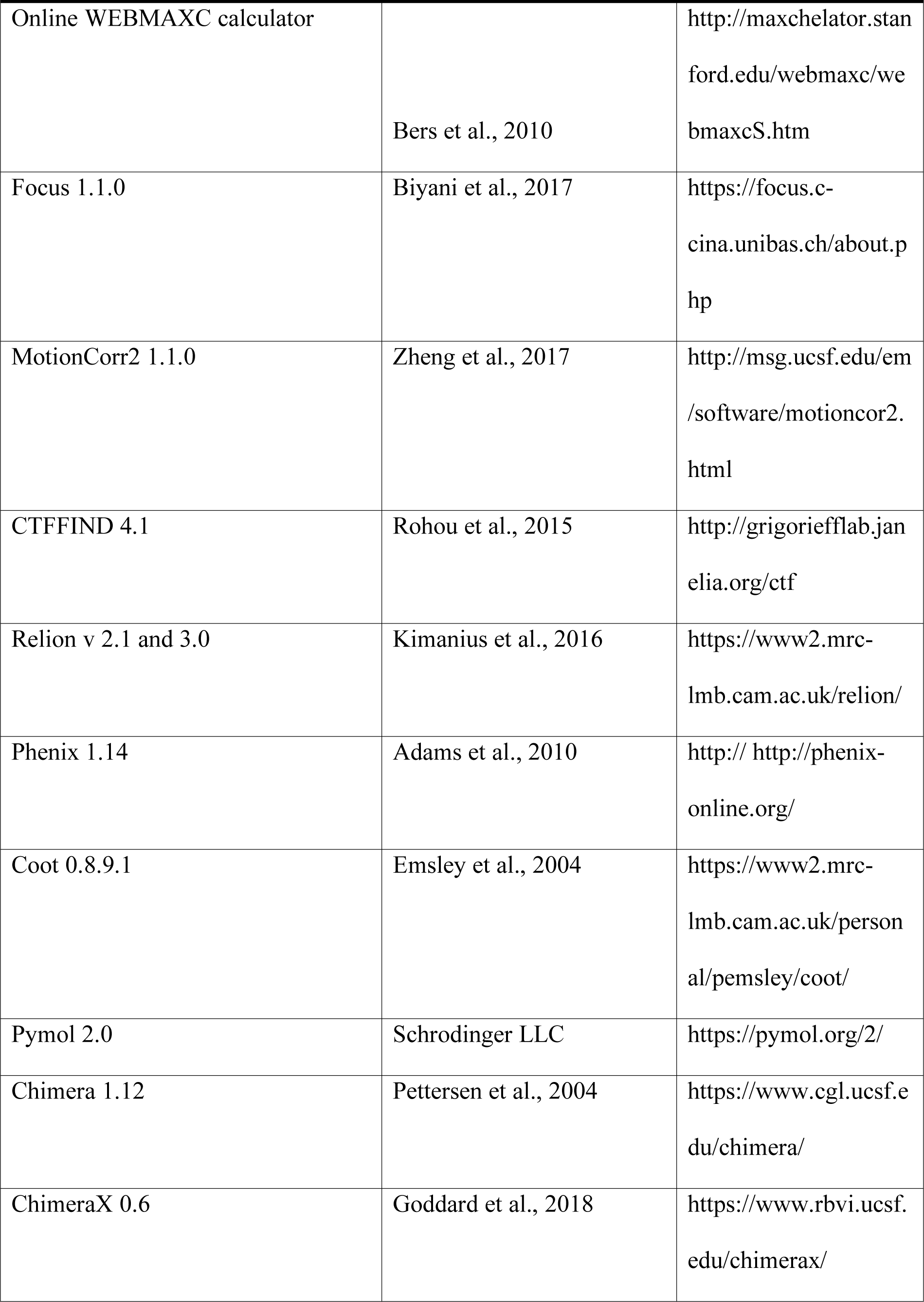

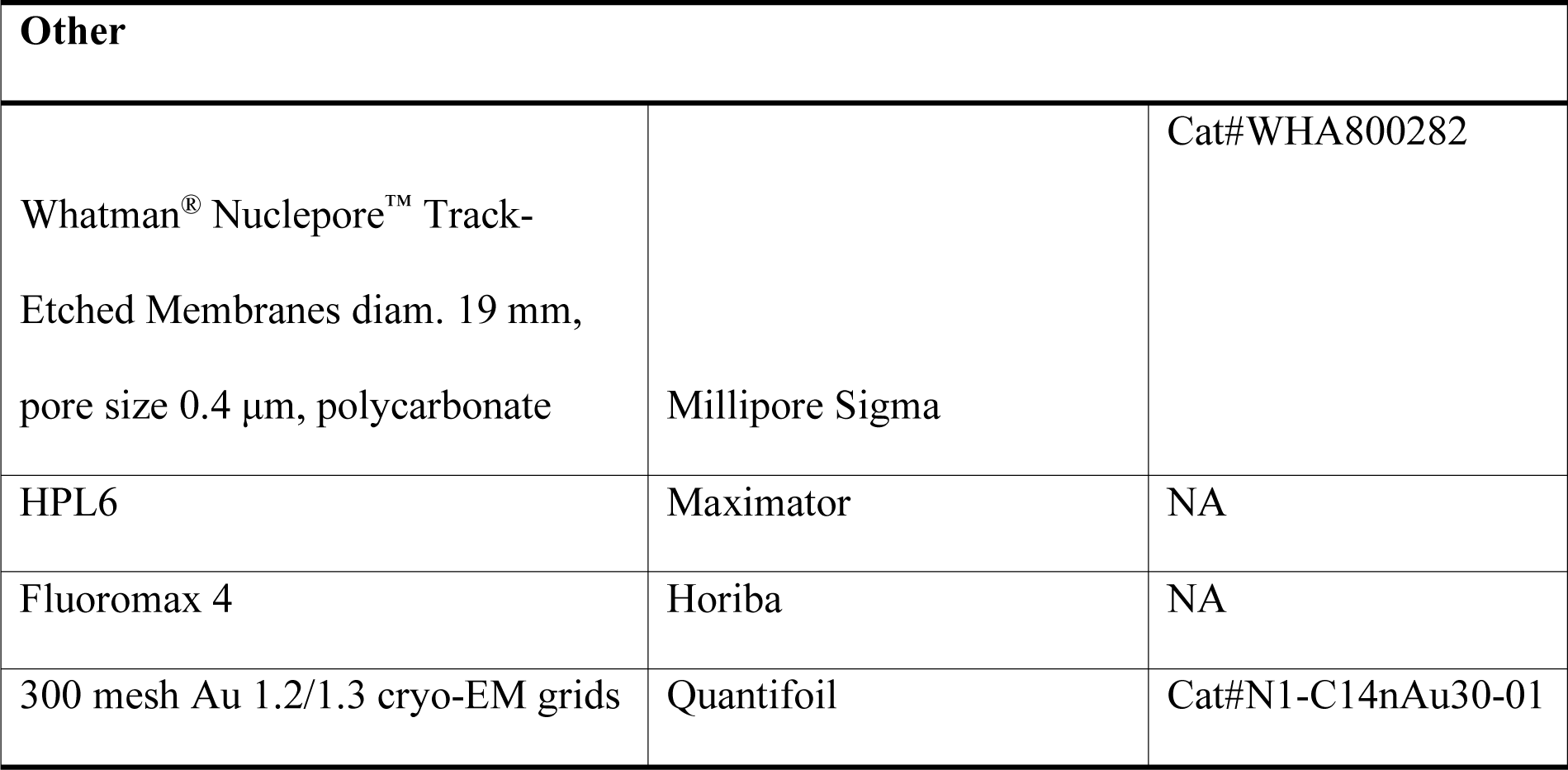

### Strains

Wild type *S. cerevisiae* FGY217 were grown either on YPD agar or in YPD liquid media supplemented with 2% glucose at 30 °C. After the transformation with respective plasmids, the cells were grown on yeast synthetic drop-out media without uracil with 2% glucose at 30 °C. For protein expression, the cells were transferred into a selective media containing 0.1% glucose.

### Construct preparation

The sequence encoding nhTMEM16 was cloned into a modified FX-cloning compatible (Geertsma and Dutzler, 2011) pYES2 vector as a C-terminal fusion to streptavidin-binding peptide (SBP) and Myc tags that were followed by an HRV 3C cleavage site. The mutations were introduced using a QuickChange method (Zheng et al., 2004).

### Protein expression and purification

All buffers were prepared using Ca^2+^-free water for molecular biology (Millipore). Plasmids carrying the WT or mutant genes were transformed into the *S. cerevisiae* FGY217 strain as described (Gietz and Schiestl, 2007) using lithium acetate/single-stranded carrier DNA/polyethylene glycol method. The cells carrying the plasmid were grown at 30 °C in a yeast synthetic dropout media until the culture reached the OD_600_ of 0.8. Afterwards, the protein expression was initialized by adding 2% galactose and the temperature was decreased to 25 °C. The protein was expressed for 40 hours after induction. Cells were harvested at 7,200 *g* for 10 min and resuspended in buffer A (50 mM HEPES pH 7.6, 150 mM NaCl, 10 mM EGTA) containing 1 mM MgCl_2_, DNAse and protease inhibitor cocktail tablets and lysed using a high pressure cell lyser HPL6 at 40 KPsi. Cell-debris was removed by centrifugation at 8,000 *g* for 30 min. Membranes were subsequently harvested by centrifugation with a 45 TI rotor (Beckmann) at 200,000 *g* for 1.5 hours, resuspended in buffer A containing 5% glycerol and flash-frozen in liquid N_2_ and stored at −80 °C until further use. During purification, all steps were carried out at 4 °C or on ice. For purification, different amount of membrane vesicles were used depending on the purpose of the experiment: 30 g of membranes were used for sample preparation for cryo EM in detergent and in nanodiscs each, and 10 g of membranes for functional assays. Membranes were solubilized in buffer A containing 2% *n***-**dodecyl**-**β**-**d**-**maltopyranoside (DDM) and 5% glycerol for 1.5 h. The insoluble fraction was removed by ultracentrifugation at 200,000 *g* for 30 min. The supernatant was applied to 4 ml of streptavidin Ultralink resin for 1.5 h, incubated under gentle agitation. The resin containing bound protein was subsequently washed with 50 column volumes (CV) of buffer B (10 mM HEPES pH 7.6, 150 mM NaCl, 5 mM EGTA, 0.03% DDM, 5% glycerol).

For sample preparation for cryo-EM in detergent, the purification tag was cleaved on the resin with 2.4 mg of HRV 3C protease in buffer B containing 20 μg/ml of yeast polar lipids for 2 h. The flow-through was collected, concentrated using 100 kDa cut-off centrifugal filter units at 700 *g* and injected onto a Superdex 200 size-exclusion column equilibrated in buffer C (5 mM HEPES pH 7.6, 150 mM NaCl, 2 mM EGTA, 0.03% DDM). Main peak fractions were pooled and concentrated to 3.3 mg/ml as described above.

For sample preparation in lipid nanodiscs, membrane scaffold protein (MSP) 2N2 was expressed and purified as described (Ritchie et al., 2009), except that the polyhistidine-tag was not removed. Chloroform-solubilized lipids (POPC:POPG at a molar ratio of 7:3) were pooled, dried under a nitrogen stream and washed twice with diethyl ether. The resulting lipid film was dried in a desiccator overnight, and solubilized in 30 mM DDM at a final lipid concentration of 10 mM. nhTMEM16 was purified as described above using the same amount of streptavidin resin, except that the uncleaved fusion construct was eluted from the beads with buffer B containing 3 mM biotin. Protein-containing fractions were pooled and concentrated as described above. Biotin was removed from the sample via gel filtration on a Superdex 200 column. Purified protein was assembled into 2N2 nanodiscs at three different molar ratios of protein:lipids:MSP (*i.e.* 2:725:10, 2:775:10, and 2:1100:10) as described (Ritchie et al., 2009). This procedure resulted in a five-fold excess of empty nanodiscs, which prevented multiple incorporations of target proteins into a single nanodisc. nhTMEM16 was mixed with detergent-solubilized lipids and incubated for 30 min on ice. Subsequently, purified 2N2 was added to the sample, and the mixture was incubated for an additional 30 min. Detergent was removed by incubating the sample overnight with SM-2 biobeads (200 mg of beads/ml of the reaction) under constant rotation. From this point on, detergent was excluded from all buffers. To separate the protein-containing from empty nanodiscs, the sample was incubated with 1 ml of streptavidin resin for 1.5 hours. The resin was washed with 10 CV of buffer B and assembled nanodiscs containing nhTMEM16 were eluted in buffer B containing 3 mM biotin. The purification tag was removed by incubation with 0.8 mg of HRV 3C protease for 2 hours. Cleaved samples were concentrated at 500 *g* using concentrators (Amicon) with a molecular weight cut-off of 100 kDa and injected onto a Superose 6 column equilibrated in buffer C. Analogously to the detergent sample, the main peak fractions were concentrated to around 2 mg/ml.

For reconstitution into proteoliposomes, nhTMEM16 was purified as described for detergent samples used for cryo-EM with minor differences: lysate was incubated with 3 ml of streptactin resin and eluted in buffer B containing 5 mM of d-desthiobiotin and the purification tag was not removed.

### Reconstitution into the liposomes and scrambling assay

Functional reconstitution was carried out essentially as described (Malvezzi et al., 2013). Lipids for the reconstitution *(E. coli* polar extract and egg-PC at a ratio of 3:1 (w/w) was supplemented with 0.5% (w/w) 18:1-06:0 NBD-PE) were pooled and dried as described above. The lipid film was dissolved in assay buffer A (20 mM HEPES pH 7.5, 300 mM KCl, 2 mM EGTA) containing 35 mM CHAPS at a final lipid concentration of 20 mg/ml, aliquoted, flash-frozen and stored until further use. Soy bean lipids extract with 20% cholesterol were prepared as described in Alvadia et al, 2018. On the day of reconstitution, lipids were diluted to 4 mg/ml in assay buffer A and purified protein was added to the lipid mixture at a lipid to protein ratio of 300:1 (w/w). As a control, an equivalent volume of the gel filtration buffer was added to aliquots from the same batch of lipids and treated the same way as the proteoliposomes sample, resulting in the assembly of empty liposomes. Soy bean lipid mixture was destabilized with Triton X-100 as described in Alvadia et al, 2018. The mixture was incubated for 15 min at room temperature (RT). Biobeads (15 mg of beads/mg of lipids) were added 4 times (after 15 min, 30 min, 1 h and 12 h). After the second addition of biobeads, the sample was transferred to 4 °C. The proteoliposomes were harvested by centrifugation (150,000 *g*, 30 min, 4 °C), resuspended in assay buffer A containing desired amounts of free Ca^2+^ (calculated with the WEBMAXC calculator) at a final concentration of 10 mg/ml in aliquots of 250 μl, flash-frozen and stored at −80 °C until further use. To characterize the protein activity in the lipid mix used for the preparation of nanodisc samples, nhTMEM16 was reconstituted into liposomes composed of 7 POPC:3 POPG (mol/mol) with 0.5 % 18:1-06:0 NBD-PE. For conditions mimicking a eukaryotic membrane, the protein was reconstituted in soybean polar lipids extract with 20% cholesterol (w/w) and 0.5% 18:1-06:0 NBD-PE. Each mutant was purified and reconstituted independently three times on three different days (defined as distinct biological replicate). To correct for differences in the reconstitution efficiency, WT was included alongside with the mutants into each experiment, and reconstituted into the same batch of lipids on the same day. Reconstitution of the WT into nanodisc lipids and soy bean lipids was performed once.

On the day of the measurement, 250 μl aliquots of liposomes were frozen and thawed 3 times and extruded 21 times through a 400 nm polycarbonate membrane using LipoFast extruder (Avestin). Scrambling data was recorded on a Horiba Fluoromax spectrometer. For the measurement, liposomes were diluted to 0.2 mg/ml in assay buffer B (80 mM HEPES pH 7.5, 300 mM KCl, 2 mM EGTA) containing corresponding amounts of free Ca^2+^. odium dithionite was added after 60 sec to a final concentration of 30 mM and the fluorescence decay was recorded for additional 340 sec. Traces with deviations in fluorescence values of more than 0.03×10^6^ counts per second within the first 60 s of the measurement or displaying uncharacteristic spikes in fluorescence decay after addition of dithionite were discarded. Each sample was measured three times (technical replicate). Data were normalized as F/Fmax.

Under ideal conditions, all liposomes should be unilammelar and the lipid composition of the two leaflets of the bilayer should be symmetric. Such scenario would result in a 50% fluorescence decrease upon addition of dithionite to the outside of empty liposomes and a full decay of the fluorescence in liposomes containing an active scramblase, which facilitates lipid flip-flop. In our experiments, we observed fluorescence decays to plateau values ranging between 44–60% in control liposomes and 10-20% in preparations of proteoliposomes containing nhTMEM16 constructs due to variabilities in liposome preparation. The incomplete decay is likely a consequence of multilamellar liposomes and due to a fraction of liposomes not containing an active scramblase in proteoliposome preparations. To account for differences in reconstitution efficiency, all protein constructs that are directly compared to each other in our study and that are shown on the same panel were reconstituted with the same batch of lipids on the same day and the analysis is restricted to a phenotypical comparison of the decay kinetics. Reconstitution efficiency of mutants as compared to wild type nhTMEM16 was estimated by Western blotting using a semi-dry transfer protocol. Protein was transferred onto a PVDF membrane (Immobilon-P) and detected using mouse anti-c-Myc antibody (Millipore Sigma) as a primary (1:5,000 dilution) and goat anti-mouse coupled to a peroxidase (Jackson Immunoresearch) as a secondary antibody (1:10,000 dilution).

### Cryo-electron microscopy sample preparation and imaging

2.5 µl of freshly purified protein at a concentration of 3.3 mg/ml when solubilized in DDMand at about 2 mg/ml when reconstituted in nanodiscs were applied on holey-carbon cryo-EM grids (Quantifoil Au R1.2/1.3, 200, 300 and 400 mesh), which were prior glow-discharged at 5 mA for 30 s. For datasets of Ca^2+^-bound protein, samples (containing 2mM EGTA) were supplemented with 2.3 mM CaCl_2_ 30 minutes before freezing, resulting in a free calcium concentration of 300 µM. The pH change in response to the addition Ca^2+^ was monitored and found negligible. Grids were blotted for 2–5 s in a Vitrobot (Mark IV, Thermo Fisher) at 10–15 °C and 100% humidity, plunge-frozen in liquid ethane and stored in liquid nitrogen until further use. Cryo-EM data were collected on a 200 keV Talos Arctica microscope (Thermo Fisher) at the University of Groningen using a post-column energy filter (Gatan) in zero-loss mode, a 20 eV slit, a 100 µm objective aperture, in an automated fashion using EPU software (Thermo Fisher) on a K2 summit detector (Gatan) in counting mode. Cryo-EM images were acquired at a pixel size of 1.012 Å (calibrated magnification of 49,407x), a defocus range from –0.5 to –2 µm, an exposure time of 9 s with a sub-frame exposure time of 150 ms (60 frames), and a total electron exposure on the specimen of about 52 electrons per Å^2^. The best regions on the grid were screened and selected with an in-house written script to calculate the ice thickness and data quality was monitored on-the-fly using the software FOCUS (Biyani et al., 2017).

### Image Processing

For the detergent dataset collected in presence of Ca^2+^, the 2,521 dose-fractionated cryo-EM images recorded (final pixel size 1.012 Å) were subjected to motion-correction and dose-weighting of frames by MotionCor2 (Zheng et al., 2017). The CTF parameters were estimated on the movie frames by ctffind4.1 (Rohou and Grigorieff, 2015). Images showing contamination, a defocus above –0.5 or below –2 µm, or a poor CTF estimation were discarded. The resulting 2,023 images were used for further analysis with the software package RELION2.1 (Kimanius et al., 2016). Around 4,000 particles were initially manually picked from a subset of the dataset and classified to create a reference for autopicking. The final round of autopicking on the whole dataset yielded 251,693 particles, which were extracted with a box size of 220 pixels and initial classification steps were performed with two-fold binned data. False positives were removed in the first round of 2D classification. Remaining particles were subjected to several rounds of 2D classification, resulting in 174,901 particles that were further sorted in several rounds of 3D classification. A map created from the X-ray structure of nhTMEM16 (PDBID: 4WIS) was low-pass filtered to 50 Å and used as initial reference for the first round of 3D classification. The resulting best output class was used as new reference in subsequent jobs in an iterative way. The best 3D classes, comprising 128,648 particles, were subjected to auto-refinement, yielding a map with a resolution of 4.3 Å. In the last refinement iteration, a mask excluding the micelle was used and the refinement was continued until convergence (focused refinement), which improved the resolution to 4.0 Å. The final map was masked and sharpened during post-processing resulting in a resolution of 3.9 Å. Finally, the newly available algorithms for CTF refinement and Bayesian polishing implemented in Relion3.0, were applied to further improve the resolution (Zivanov et al., 2018). A final round of 3D classification was performed, resulting in 120,086 particles that were subjected to refinement, providing a mask generated from the final PDB model in the last iteration. The final map at 3.6 Å resolution was sharpened using an isotropic B-factor of −126 Å^2^. While a C1 symmetry was tested and applied throughout image processing workflow, no indication of an asymmetry was identified. Therefore, a C2-symmetry was imposed during the final 3D classification and auto-refinement. Local resolution was estimated by RELION. All resolutions were estimated using the 0.143 cut-off criterion (Rosenthal and Henderson, 2003) with gold-standard Fourier shell correlation (FSC) between two independently refined half maps (Scheres and Chen, 2012). During post-processing, the approach of high-resolution noise substitution was used to correct for convolution effects of real-space masking on the FSC curve (Chen et al., 2013). The directional resolution anisotropy of density maps was quantitatively evaluated using 3DFSC (Tan et al., 2017).

For the other datasets, a similar workflow for image processing was applied. In case of the detergent dataset collected in absence of Ca^2+^, a total of 570,203 particles were extracted with a box size of 240 pixels from 2,947 images. Several rounds of 2D and 3D classification resulted in a final number of 295,219 particles, which yielded a 3.9 Å map after refinement and post-processing. CTF refinement and Bayesian polishing were applied, followed by a second round of CTF refinement. One last round of 3D classification was performed, leading to a final pool of 238,070 particles which, after refinement and post processing, resulted in a map at 3.7 Å resolution. For structure determination of nhTMEM16 in lipid nanodiscs, small datasets (around 1000 images) recorded from samples reconstituted at different lipid to protein ratios (LPR) in absence of Ca^2+^ during nanodisc assembly, identified the lowest LPR as optimal condition for large-scale data collection. The dataset in nanodisc in absence of Ca^2+^ resulted in 1,379,187 auto-picked particles from 6,465 images extracted with a box size of 240 pixels, which were reduced to 150,421 particles after several rounds of 2D and 3D classification. CTF refinement and Bayesian polishing followed by a final round of 3D classification was performed, resulting in a selection of 133,961 particles. The final refinement and masking resulted in a 3.8 Å resolution map.

Finally, for the dataset in nanodiscs in the presence of Ca^2+^, 2,440,110 particles were picked from 9,426 images and extracted with a box size of 240 pixels. Since the dataset displayed high conformational heterogeneity, C1 symmetry was applied throughout the processing workflow. After several rounds of 2D and 3D classification, the best classes (319,530 particles) were combined and refined. The resulting map was used for CTF refinement in Relion3. Refined particles were subjected to 3D refinement and the obtained map was used to generate a mask for nanodisc signal subtraction. The resulting subtracted particles were used for another round of 3D classification, yielding a subset of 279,609 particles. It was evident from the processing results that the core of the protein was stable and displayed a single conformation, whereas α-helices 3 and 4 appeared to be mobile. To separate distinct conformational states, all the particles were aligned using 3D refinement and subsequently subjected to a 3D classification without alignment, using the angles obtained from the preceding refinement job. In one of the resulting classes, the density corresponding to helices 3, 4 and 6 of one of the monomers was not defined, and these particles (11,25%) were thus excluded from further processing. Of the remaining particles, 36,5% are heterogeneous in the region corresponding to α-helices 3 and 4, 14,89% represent nhTMEM16 in a closed conformation of the subunit cavity, 11,91% in the intermediate, and two classes (consisting of 15,64% and 9,81% particles respectively) display an open conformation. Each class was subsequently initially refined applying C1 symmetry. Since no difference between the conformations of individual subunits in the dimeric protein was detected, the refinement was repeated with C2 symmetry imposed. The two open classes were merged and refined together. Similar to other datasets, masks generated from PDB models were applied in the last iteration of the refinement. After refinement and post-processing, the resulting resolutions were 3.57 Å for the open class, 3.57 Å for the closed class and 3.68 Å for the intermediate class. We also attempted to classify the heterogeneous class further, but could not improve the separation of the particles. To generate the map of the open conformation containing the nanodisc signal, we reverted the particles from the open class to their original non-subtracted equivalent, and refined them applying C2 symmetry. This map was used to prepare Figure 3-supplement 3, Figure 6 and Video 2.

### Model building refinement and validation

For the detergent dataset in the presence of Ca^2+^, the X-ray structure of nhTMEM16 (PDBID: 4WIS) was used as a template. The model was manually edited in Coot (Emsley and Cowtan, 2004) prior to real-space refinement in Phenix imposing symmetry restraints between subunits throughout refinement (Adams et al., 2010). The higher quality of the cryo-EM density of the cytoplasmic domains compared to the X-ray density allowed for the identification of an incorrect register at the N-terminus of the X-ray structure (PDBID: 4WIS) between residues 20-61. The region comprising residues 15-61 was thus rebuilt. Another change compared to the X-ray structure concerns the remodeled conformation of Lys 598. In several cases the cryo-EM density allowed the interpretation of residues that were not defined in the X-ray structure. These concern residues 130-140 at a loop connecting the cytoplasmic and the transmembrane domain, an extra helical turn at the cytoplasmic side of α6 and residues 476-482 at the loop connecting α6 to α6’. Since residues 690-698 appeared to be helical in the cryo-EM map, the region was rebuilt and it was possible to add the missing residues 688-691. Several regions of the cryo-EM map displayed weaker density compared to the X-ray structure, and hence the residues 416-418, 653-656 and 660-663 had to be removed from the model. This corrected nhTMEM16 structure (PDBID: 6QM5) was used for refinement of the nhTMEM16 structure in DDM in the absence of Ca^2+^, and for model building and refinement of the Ca^2+^-bound open and Ca^2+^-bound closed structures in nanodiscs. All models were edited manually in Coot, followed by real space refinement in Phenix. The Ca^2+^-free structure in DDM (PDBID: 6QM6) and the Ca^2+^-bound open structure in nanodisc (PDBID: 6QM9) closely resemble the Ca^2+^-bound structure in DDM, displaying only minor differences. Since the density corresponding to α6 is better resolved in the Ca^2+^-bound open structure in nanodisc, the helix was extended to Ser 471. The density corresponding to Cα2 of the C-terminus (residues 664-684) was comparably weak in all three open structures and the helix was thus fitted as a rigid body. For the Ca^2+^-bound closed conformation (PDBID: 6QMB), major changes were introduced for α3, α4 and to a lesser extent for α6. The densities of the loop connecting α5’ and α6 and of the cytosolic region of α6 were generally weaker and residues 417-424 and 467-471 were thus not interpreted in this structure. Since the Ca^2+^-bound closed map in nanodisc displays the highest resolution and quality in the region encompassing α3 and α4, the resulting model was used as a template for the interpretation of the Ca^2+^-bound intermediate (PDBID: 6QMA) and Ca^2+^-free (PDBID: 6QM4) structures in nanodisc. In case of the Ca^2+^-bound intermediate structure, introduced changes are mostly confined to α4, where the extracellular part of the α-helix undergoes a conformational transition towards opening of the subunit cavity. For the Ca^2+^-free structure in nanodisc, the residues 461-467 of α6 were not modelled due to its high mobility. Figures were prepared using Pymol (The PyMOL Molecular Graphics System, Version 2.0 Schrödinger, LLC), Chimera (Pettersen et al., 2004) and ChimeraX (Goddard et al., 2018).

## Acknowledgments

We thank S. Klauser, S. Rast and M. Punter for their help in establishing the computer infrastructure, H. Stahlberg and K. Goldie at C-Cina of the University of Basel for access to cryo-electron microscopes at an initial stage of the project. S. Weidner and the Workshop of Biochemistry department (UZH) are acknowledged for developing the high-pressure cell lyser. D. Deneka is acknowledged for his advice on nanodisc reconstitution. All members of the Dutzler and Paulino labs are acknowledged for help at various stages of the project.

## Additional Information

### Funding

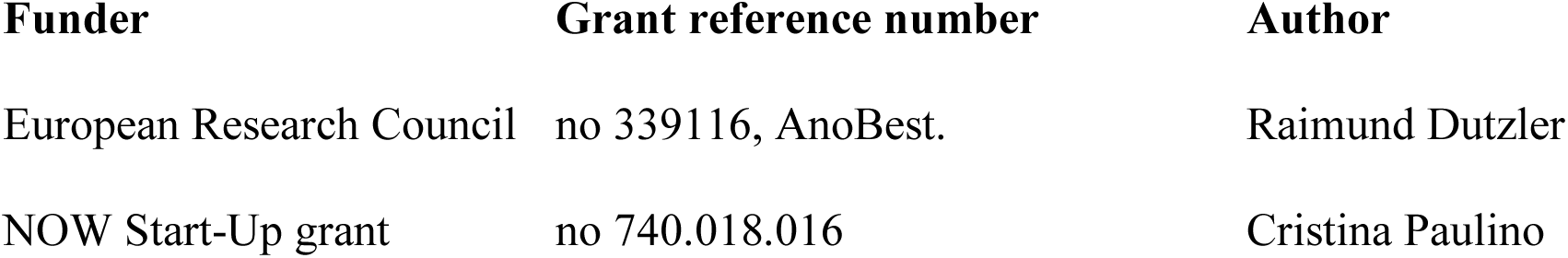

The funders had no role in study design, data collection and interpretation, or the decision to submit the work for publication.

## Author contributions

V.K. and L.B. purified proteins for cryo-EM and functional characterization. V.K. reconstituted protein into nanodiscs and liposomes and carried out lipid transport experiments. V.K. and C.P. prepared the samples for cryo-EM. V.K., V.C.M., G.T.O. and C.P. collected cryo-EM data. V.K., V.C.M and C.P. carried out image processing, model building and refinement. V.K., V.C.M, R.D. and C.P. jointly planned experiments, analyzed the data and wrote the manuscript.

## Additional files

### Data availability

The three-dimensional cryo-EM density maps of calcium-free nhTMEM16 in detergent and nanodiscs have been deposited in the Electron Microscopy Data Bank under accession numbers EMD-4589 and EMD-4587, respectively. The maps of calcium-bound samples in detergent and calcium-bound open, calcium-bound intermediate and calcium-bound closed in nanodiscs were deposited under accession numbers EMD-4588, EMD-4592, EMD-4593 and EMD-4594, respectively. The deposition includes the cryo-EM maps, both half-maps, the unmasked and unsharpened refined maps and the mask used for final FSC calculation. Coordinates of all models have been deposited in the Protein Data Bank. The accession numbers for calcium-bound and calcium-free models in detergents are 6QM5 and 6QM6, respectively. The accession numbers for calcium-bound open, calcium-bound intermediate, calcium-bound closed and calcium-free models in nanodiscs are 6QM9, 6QMA, 6QMB and 6QM4, respectively.

## Figures

**Figure 1-figure supplement 1.**
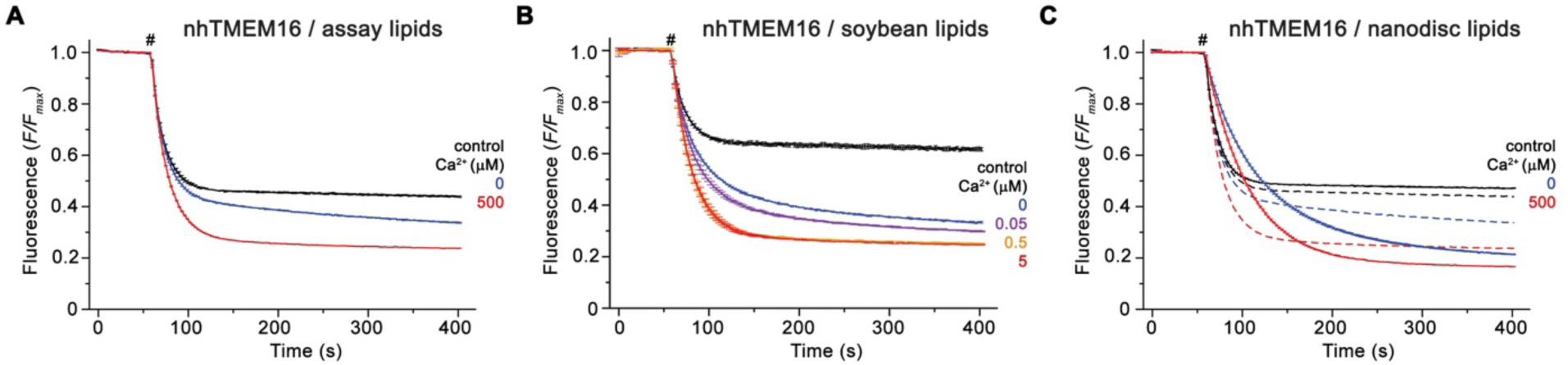
Reconstitution of nhTMEM16 into liposomes. Ca^2+^-dependence of scrambling activity of WT in liposomes of three different lipid compositions. Traces depict the fluorescence decrease of tail-labeled NBD-PE lipids after addition of dithionite (#) at different Ca^2+^ concentrations. Data show averages of three technical replicates for assay and nanodisc lipids, and of two for soybean lipids. Ca^2+^ concentrations (µM) are indicated. (**A**) Scrambling activity of nhTMEM16 reconstituted into liposomes composed of *E. coli* polar lipids, egg PC at a ratio of 3:1 (w/w) used for functional experiments. (**B**) Scrambling activity of nhTMEM16 reconstituted into liposomes composed of soybean polar lipids extract with 20% cholesterol (w/w). (**C**) Scrambling activity of nhTMEM16 reconstituted into liposomes composed of POPC/POPG at a molar ratio of 7:3 used for nanodisc assembly. Traces of *E. coli* polar lipids/egg PC-containing proteoliposomes (displayed in A) are shown as dashed lines for comparison.

**Figure 1-figure supplement 2.**
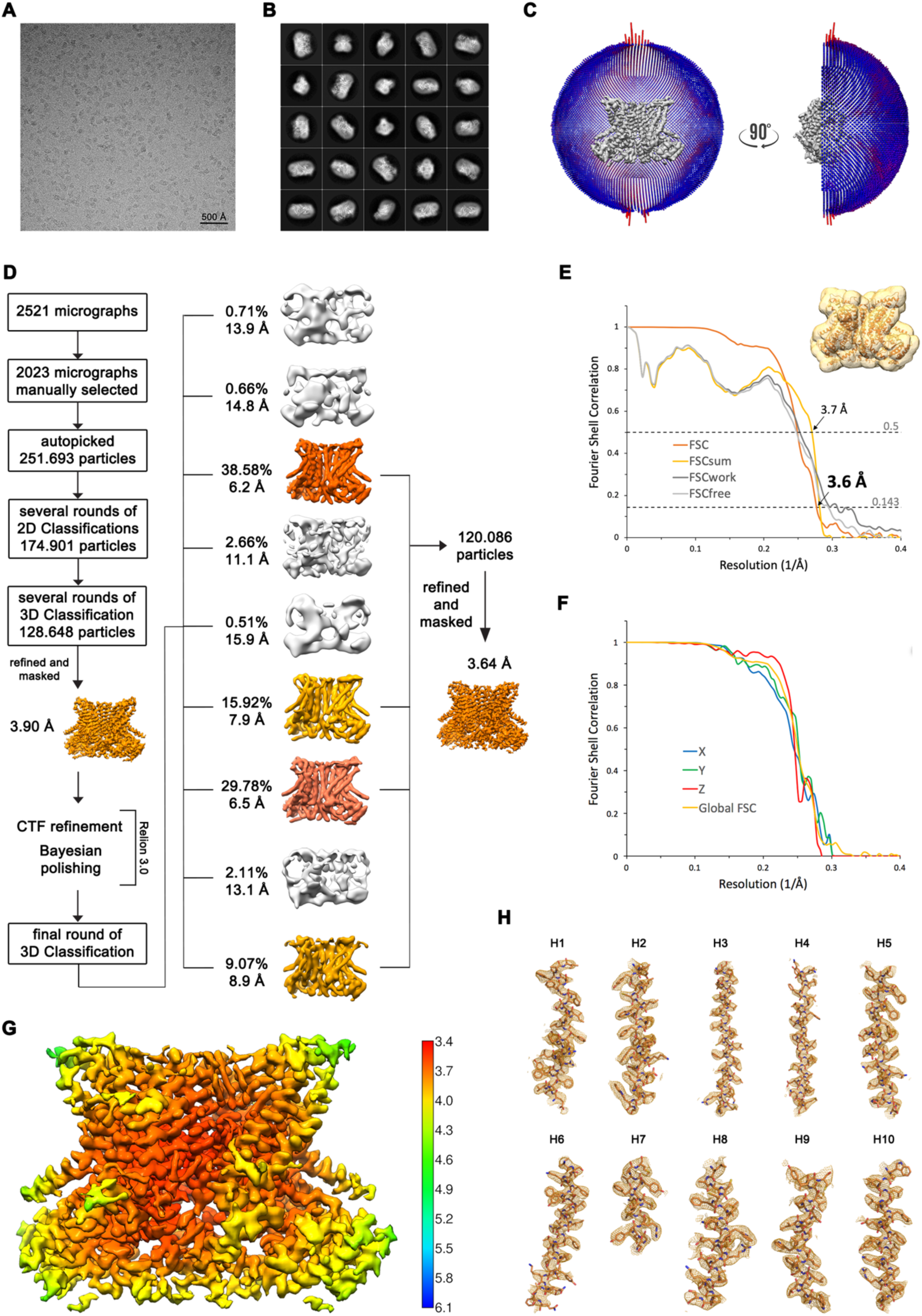
Structure Determination of nhTMEM16 in DDM in complex with Ca^2+^. (**A**) Representative cryo-EM image and (**B**) 2D-class averages of vitrified nhTMEM16 in a Ca^2+^-bound state in detergent. (**C**) Angular distribution plot of particles included in the final C2-symmetric 3D reconstruction. The number of particles with their respective orientation is represented by length and color of the cylinders. (**D**) Image processing workflow. (**E**) FSC plot used for resolution estimation and model validation. The gold-standard FSC plot between two separately refined half-maps is shown in orange and indicates a final resolution of 3.6 Å. FSC validation curves for FSCsum, FSCwork and FSCfree as described in the Methods are shown in light yellow, dark grey and light grey, respectively. A thumbnail of the mask used for FSC calculation overlaid on the atomic model is shown in the upper right corner and thresholds used for FSCsum of 0.5 and for FSC of 0.143 are shown as dashed lines. (**F**) Anisotropy estimation plot of the final map. The global FSC curve is represented in yellow. The directional FSCs along the x, y and z axis displayed in blue, green and red, respectively, are indicative for an isotropic dataset. (**G**) Final reconstruction map colored by local resolution as estimated by Relion, indicate regions of higher resolution. (**H**) Sections of the cryo-EM density of the map superimposed on the refined model. The model is shown as sticks and structural elements are labelled. The map was sharpened with a b-factor of –126 Å^2^ and contoured at 5 σ.

**Figure 1-figure supplement 3.**
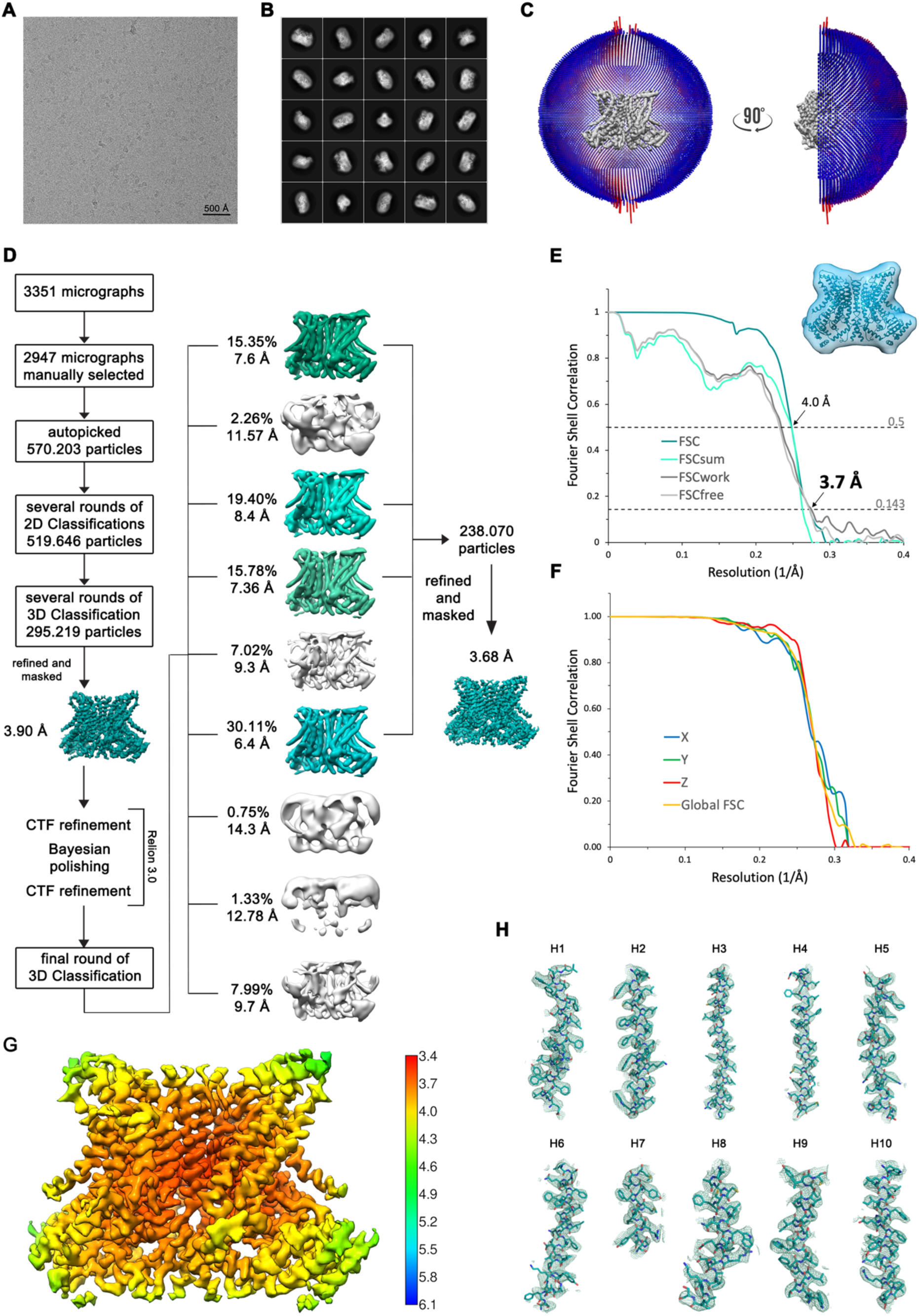
Structure Determination of nhTMEM16 in DDM in absence of Ca^2+^. (**A**) Representative cryo-EM image and (**B**) 2D-class averages of vitrified nhTMEM16 in a Ca^2+^-free state in detergent. (**C**) Angular distribution plot of particles included in the final C2-symmetric 3D reconstruction. The number of particles with their respective orientation is represented by length and color of the cylinders. (**D**) Image processing workflow. (**E**) FSC plot used for resolution estimation and model validation. The gold-standard FSC plot between two separately refined half-maps is shown in green and indicates a final resolution of 3.7 Å. FSC validation curves for FSCsum, FSCwork and FSCfree as described in the Methods are shown in cyan, dark grey and light grey, respectively. A thumbnail of the mask used for FSC calculation overlaid on the atomic model is shown in the upper right corner and thresholds used for FSCsum of 0.5 and for FSC of 0.143 are shown as dashed lines. (**F**) Anisotropy estimation plot of the final map. The global FSC curve is represented in yellow. The directional FSCs along the x, y and z axis displayed in blue, green and red, respectively, are indicative for an isotropic dataset. (**G**) Final reconstruction map colored by local resolution as estimated by Relion, indicate regions of higher resolution. (**H**) Sections of the cryo-EM density of the map superimposed on the refined model. The model is shown as sticks and structural elements are labelled. The map was sharpened with a Fig1_S4b-factor of –147 Å^2^ and contoured at 5 σ.

**Figure 1-figure supplement 4.**
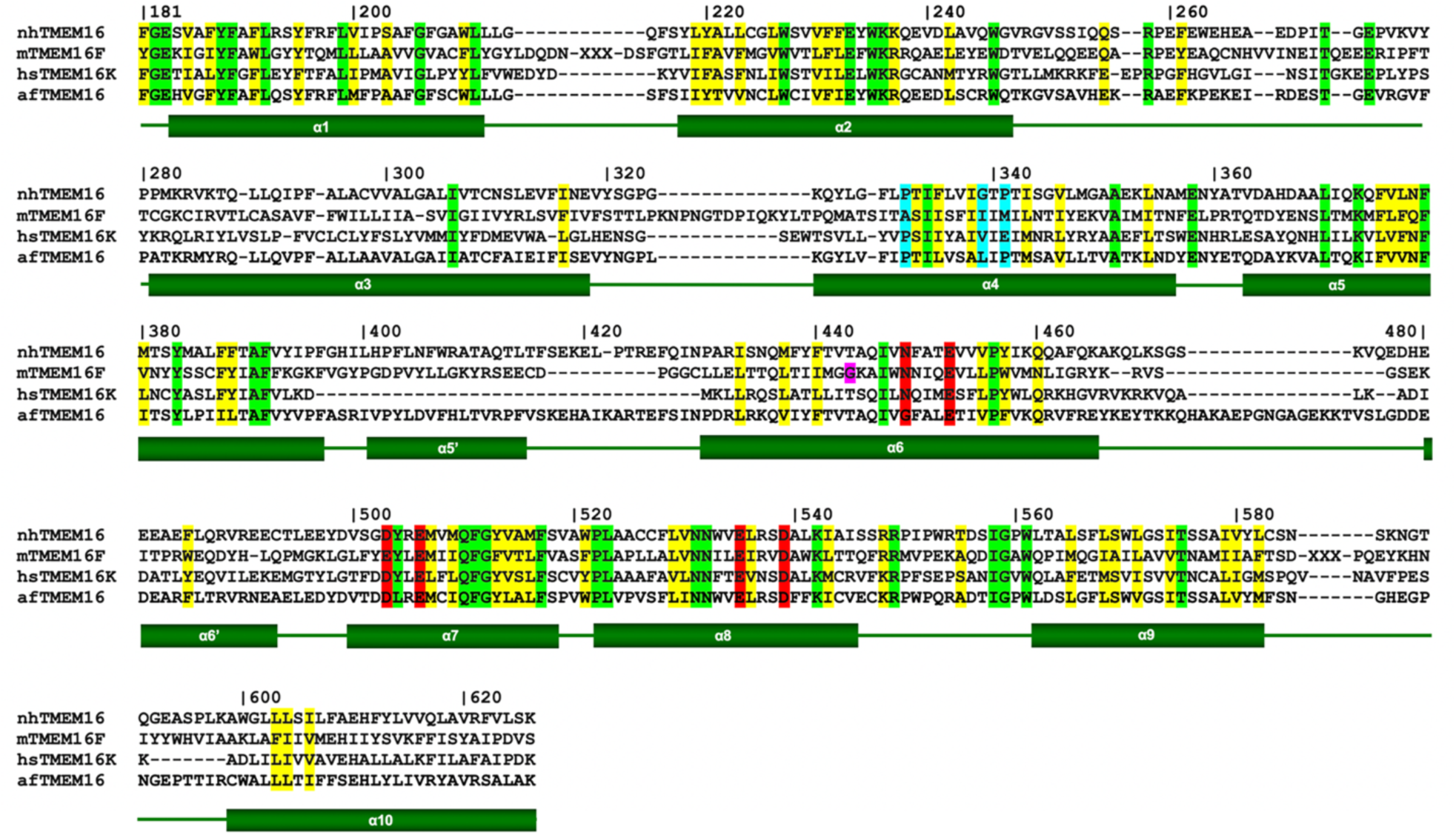
Sequence alignment of selected TMEM16 scramblases. Protein sequences of the transmembrane domain of the fungal TMEM16 homologs from *Nectria haematococca* (nhTMEM16) and *Aspergillus fumigatus* (afTMEM16), murine TMEM16F and human TMEM16K were aligned based on their structure. Numbering and the boundaries of secondary structure elements (green rectangles for transmembrane-helices and green lines for loops) correspond to nhTMEM16. Residues highlighted in red constitute the Ca^2+^-binding site; residues highlighted in blue act as potential pivot points for structural arrangements in α4; residue highlighted in purple indicates the flexible hinge region in TMEM16F responsible for the movements of α6 upon Ca^2+^ release.

**Figure 1-figure supplement 5.**
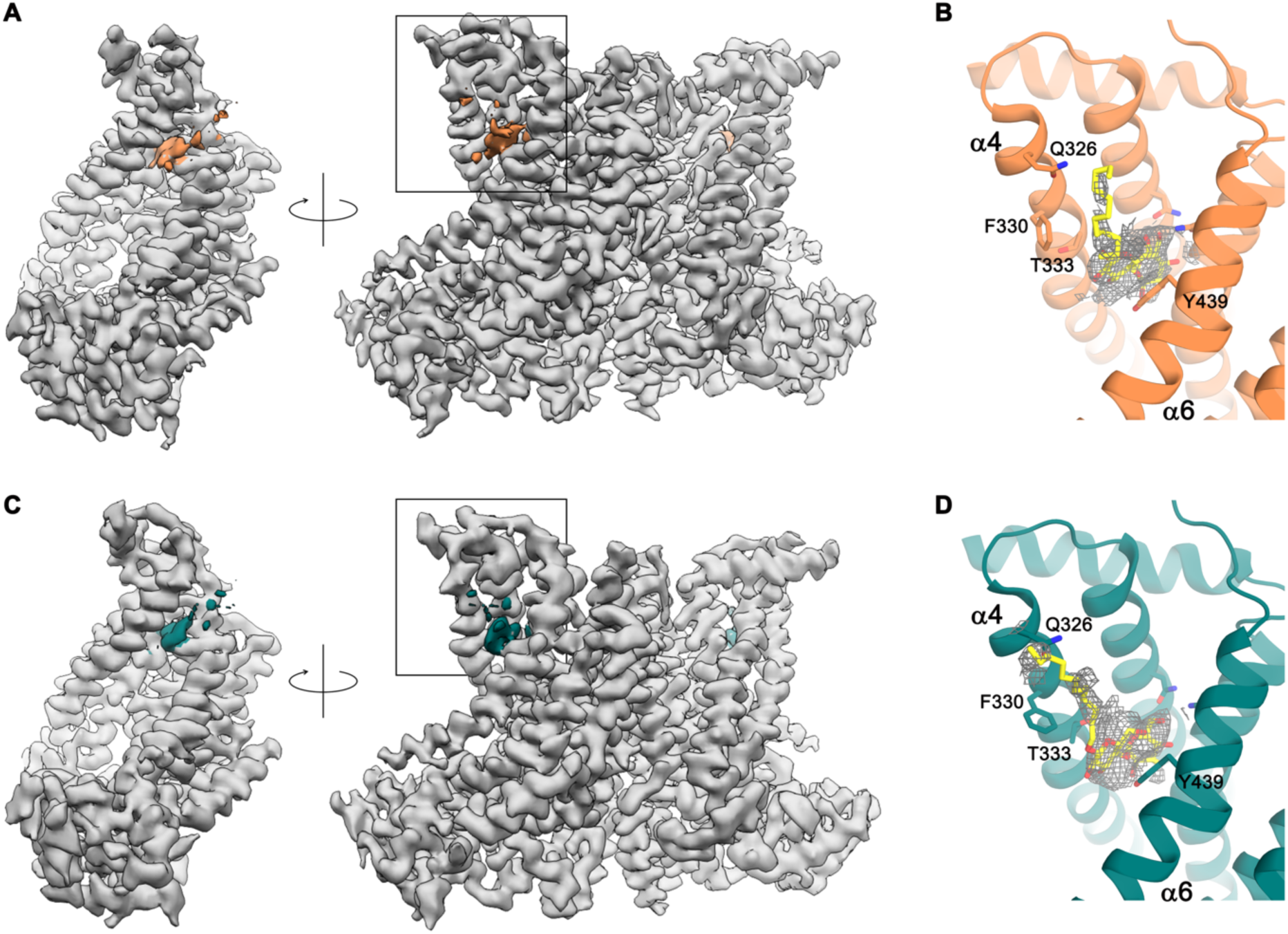
Detergent binding to the subunit cavity. (**A**) Cryo-EM map of the nhTMEM16 dimer in detergents in presence of Ca^2+^ (grey) in two orientations. Residual density in the subunit cavity highlighted in orange. (**B**) Ribbon representation of the subunit cavity of nhTMEM16 in detergents in presence of Ca^2+^. Residual density (contoured at 5 σ) is shown as grey mesh. A DDM molecule (yellow) and selected sidechains are depicted as sticks. (**C**) Cryo-EM map of the nhTMEM16 dimer in detergents in absence of Ca^2+^ (grey) in two orientations. Residual density found in the subunit cavity is highlighted in green. (**D**) Ribbon representation of the subunit cavity of nhTMEM16 in detergents in absence of Ca^2+^ (green). Residual density (contoured at 5 σ) is shown as grey mesh. A DDM molecule (yellow) is depicted as sticks. The location of B,D is indicated by the box drawn on the right panel of A,C.

**Figure 2-figure supplement 1.**
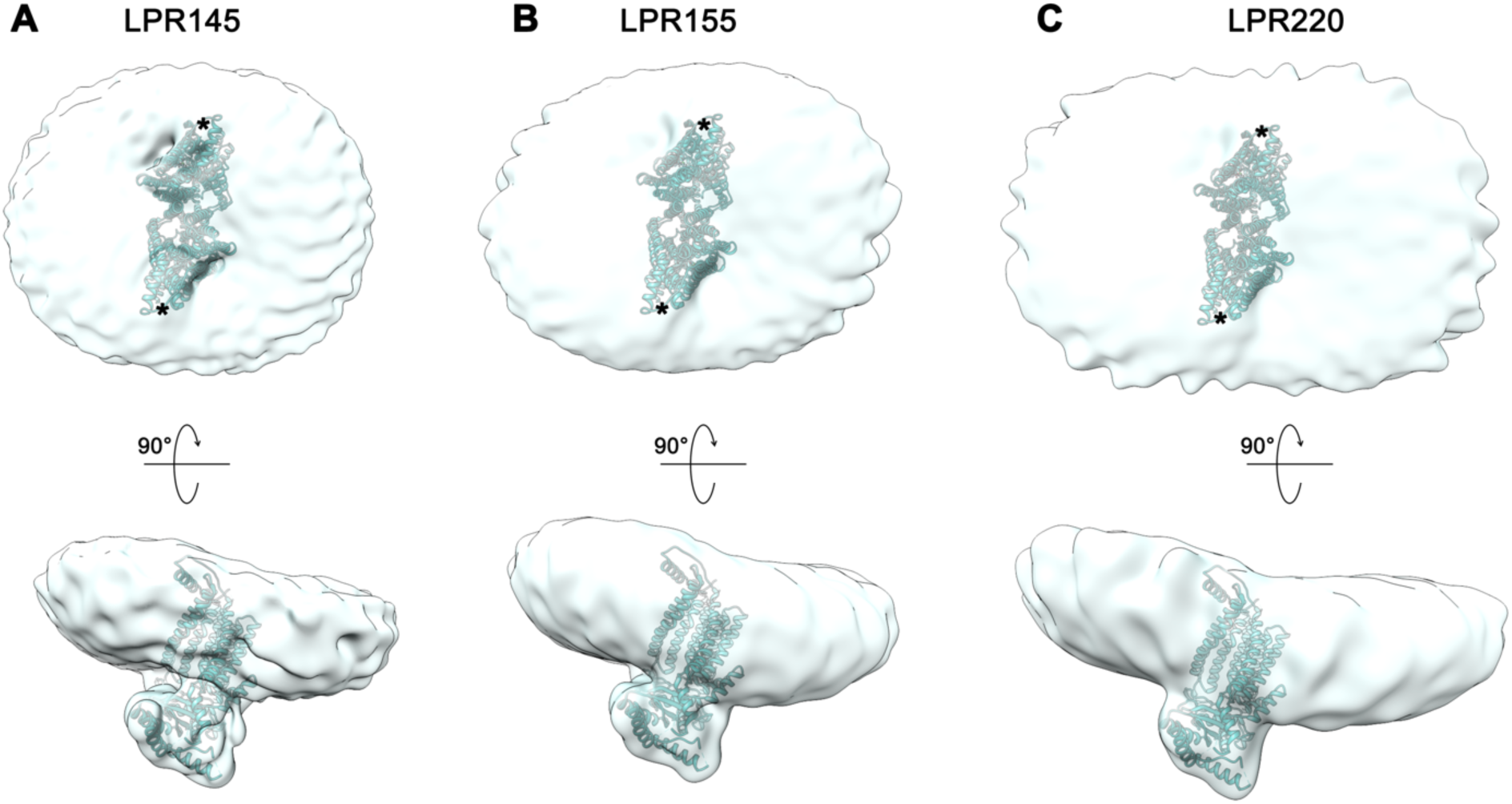
Reconstitution of nhTMEM16 into nanodiscs. Shape of nhTMEM16 particles reconstituted into nanodiscs at different protein to lipid ratios (LPR, mol/mol) in the absence of Ca^2+^: (**A**), LPR 145, (**B**), LPR 155 and(**C**), LPR 220. Refined and unmasked cryo-EM density maps were low-pass filtered to 10 Å and are shown from the extracellular side (upper panel, subunit cavity indicated by *) and from the side with a view towards the subunit cavity (lower panel). A model of nhTMEM16 fitted into the density is displayed as ribbon.

**Figure 2-figure supplement 2.**
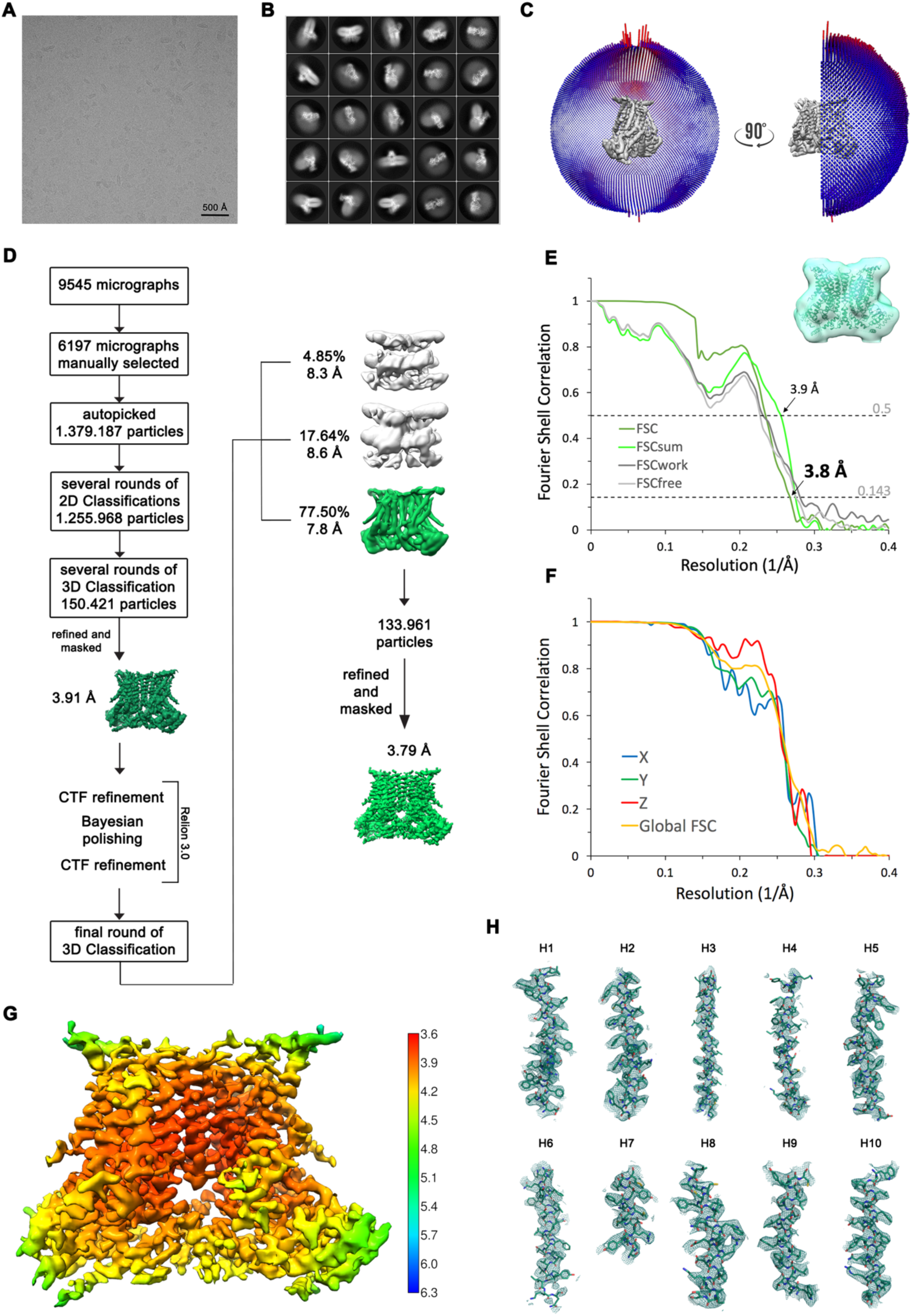
Structure Determination of nhTMEM16 in the absence of Ca^2+^ in nanodiscs. (**A**) Representative cryo-EM image and (**B**) 2D-class averages of vitrified nhTMEM16 in a Ca^2+^-free state in lipid nanodiscs. (**C**) Angular distribution plot of particles included in the final C2-symmetric 3D reconstruction. The number of particles with their respective orientation is represented by length and color of the cylinders. (**D**) Image processing workflow. (**E**) FSC plot used for resolution estimation and model validation. The gold-standard FSC plot between two separately refined half-maps is shown in green and indicates a final resolution of 3.8 Å. FSC validation curves for FSCsum, FSCwork and FSCfree as described in the Methods are shown in light green, dark grey and light grey, respectively. A thumbnail of the mask used for FSC calculation overlaid on the atomic model is shown in the upper right corner and thresholds used for FSCsum of 0.5 and for FSC of 0.143 are shown as dashed lines. (**F**) Anisotropy estimation plot of the final map. The global FSC curve is represented in yellow. The directional FSCs along the x, y and z axis displayed in blue, green and red, respectively, are indicative for an isotropic dataset. (**G**) Final reconstruction map colored by local resolution as estimated by Relion, indicate regions of higher resolution. (**H**) Sections of the cryo-EM density of the map superimposed on the refined model. The model is shown as sticks and structural elements are labelled. The map was sharpened with a b-factor of –150 Å^2^ and contoured at 5 σ.

**Figure 3-figure supplement 1.**
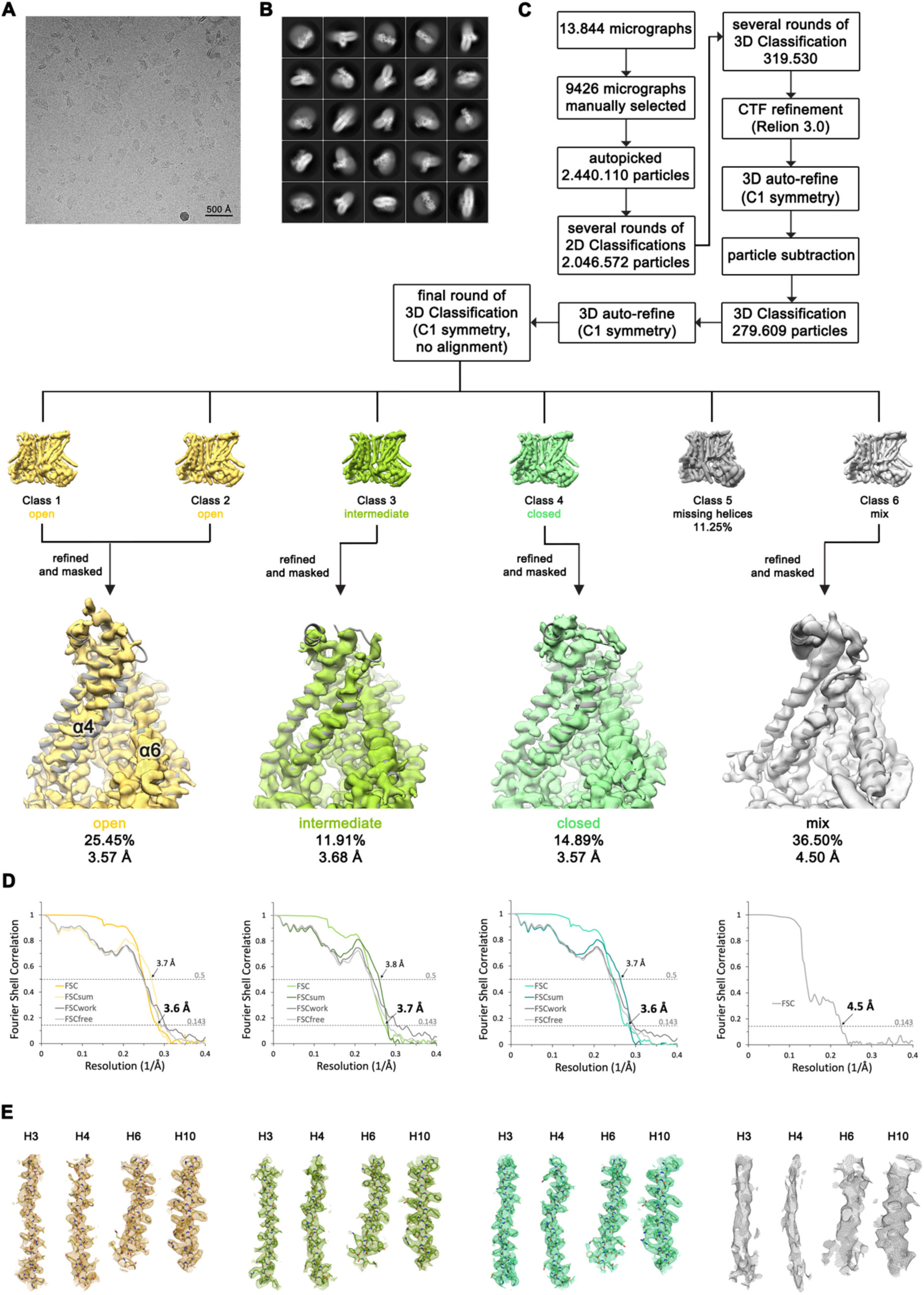
Structure Determination of nhTMEM16 in complex with Ca^2+^ in nanodiscs. (**A**) Representative cryo-EM image and (**B**) 2D-class averages of vitrified nhTMEM16 in a Ca^2+^-bound state in lipid nanodiscs. (**C**) Image processing workflow. The final set of subtracted particles was subjected to a final round of 3D classification disclosing a conformational heterogeneity at the subunit cavity, with classes resembling a well-resolved open, intermediate and closed state of the subunit cavity. Other classes displayed a poorly resolved density for α3 and α4 that likely represents an average of several transition states. Close-up of the subunit cavity show the intermediate model (grey) superimposed on the cryo-EM maps of the respective classes. View as in Figure 3C-F. (**D**) FSC plot used for resolution estimation and model validation for each state. The gold-standard FSC plot between two separately refined half-maps is shown in yellow. FSC validation curves for FSCsum, FSCwork and FSCfree as described in the Methods are shown in light yellow, dark grey and light grey, respectively. Thresholds used for FSCsum of 0.5 and for FSC of 0.143 are shown as dashed lines. (**E**) Sections of the cryo-EM density of the maps superimposed on the corresponding refined models. The models are shown as sticks and structural elements are labelled. The maps were contoured at 4.5 σ.

**Figure 3-figure supplement 2.**
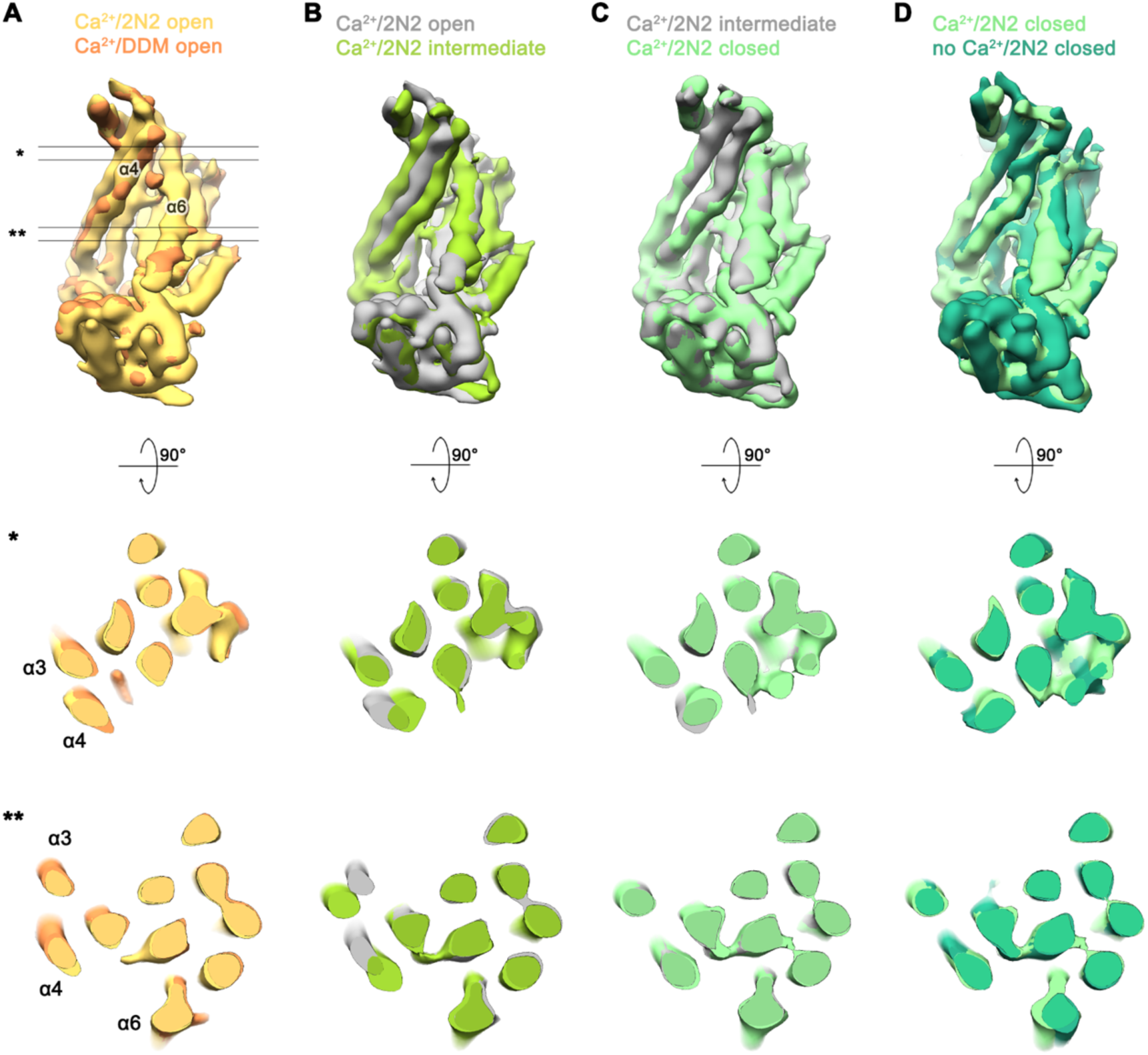
Conformational heterogeneity in the Ca^2+^-bound data of nhTMEM16 in nanodiscs. **(A-D)** Superpositions of the refined and unmasked cryo-EM maps of Ca^2+^-bound structures of nhTMEM16 in nanodiscs, low-pass filtered at 6 Å. (**A**) Superposition of the Ca^2+^-bound open structure in nanodiscs and the Ca^2+^-bound structure in DDM; (**B**) the Ca^2+^-bound open structure in nanodiscs and the Ca^2+^-bound intermediate structure in nanodiscs; (**C**) the Ca^2+^-bound intermediate structure in nanodiscs and the Ca^2+^-bound closed structure in nanodiscs; (**D**) the Ca^2+^-bound closed structure in nanodiscs and the Ca^2+^-free structure in nanodiscs. A-D, selected helices are labeled, view of upper panel is as in Figure 3C-F. Middle and lower panel show a cross-section through the superposition at the indicated positions indicated in A by * and **.

**Figure 3-figure supplement 3.**
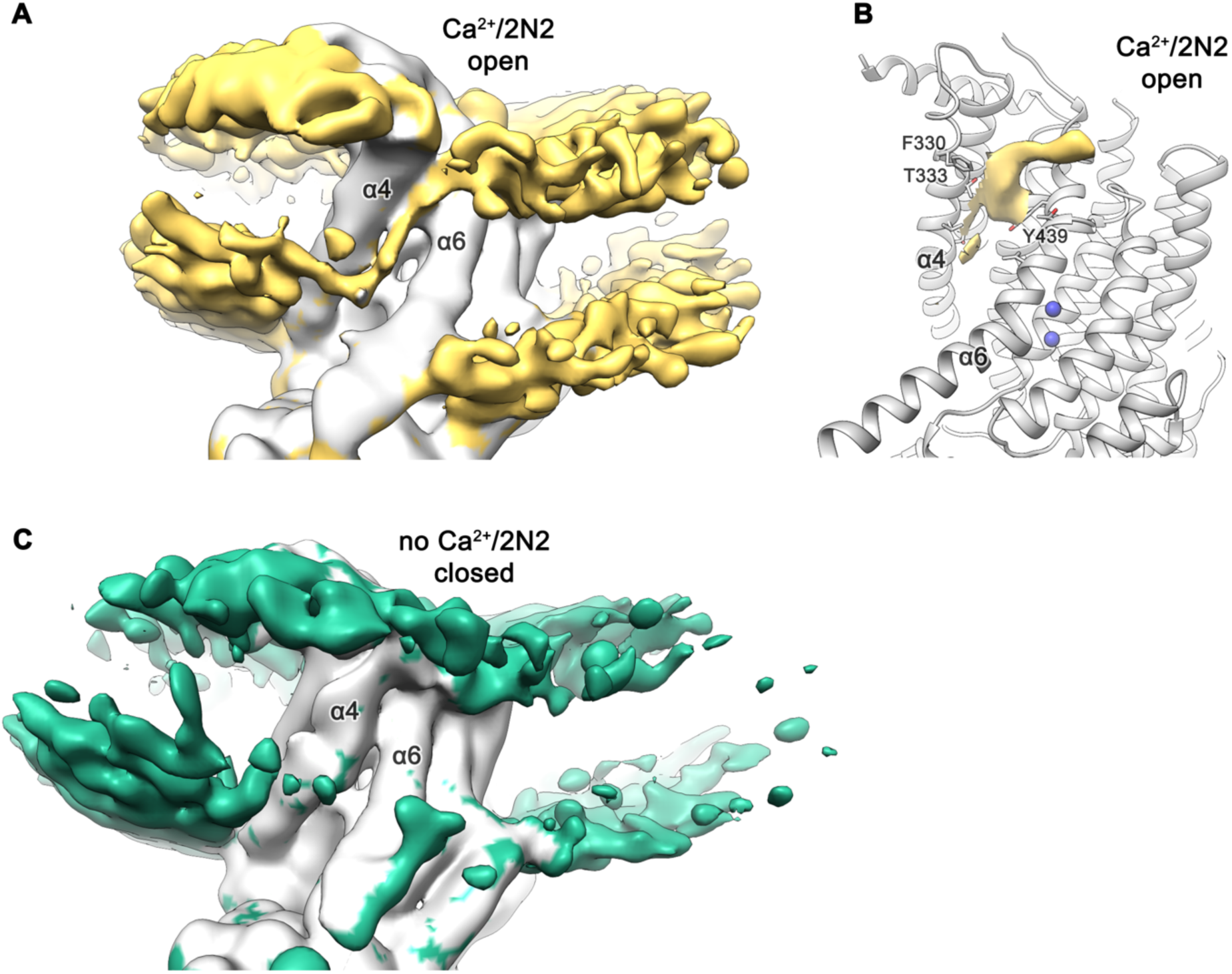
Lipid distribution at the subunit cavity. (**A**) Cryo-EM map of Ca^2+^-bound nhTMEM16 in nanodiscs (grey, contoured at 3.4 σ) with density corresponding to the phospholipid headgroup region of both membrane leaflets in the nanodisc and inside the subunit cavity highlighted in yellow. (**B**) Ribbon representation of the subunit cavity of the Ca^2+^-bound nhTMEM16 in nanodiscs (grey) with selected sidechains labelled and depicted as sticks. Unassigned density inside the subunit cavity (contoured at 3.9 σ) is shown in yellow. (**C**) Cryo-EM map of Ca^2+^-free nhTMEM16 in nanodiscs (grey, contoured at 3.4 σ) with density of the phospholipid headgroup region of both membrane leaflets in the nanodisc highlighted in green. No residual density is found at the subunit cavity. A,C The refined and unmasked maps were low-pass filtered to 6 Å.

**Figure 5-figure supplement 1.**
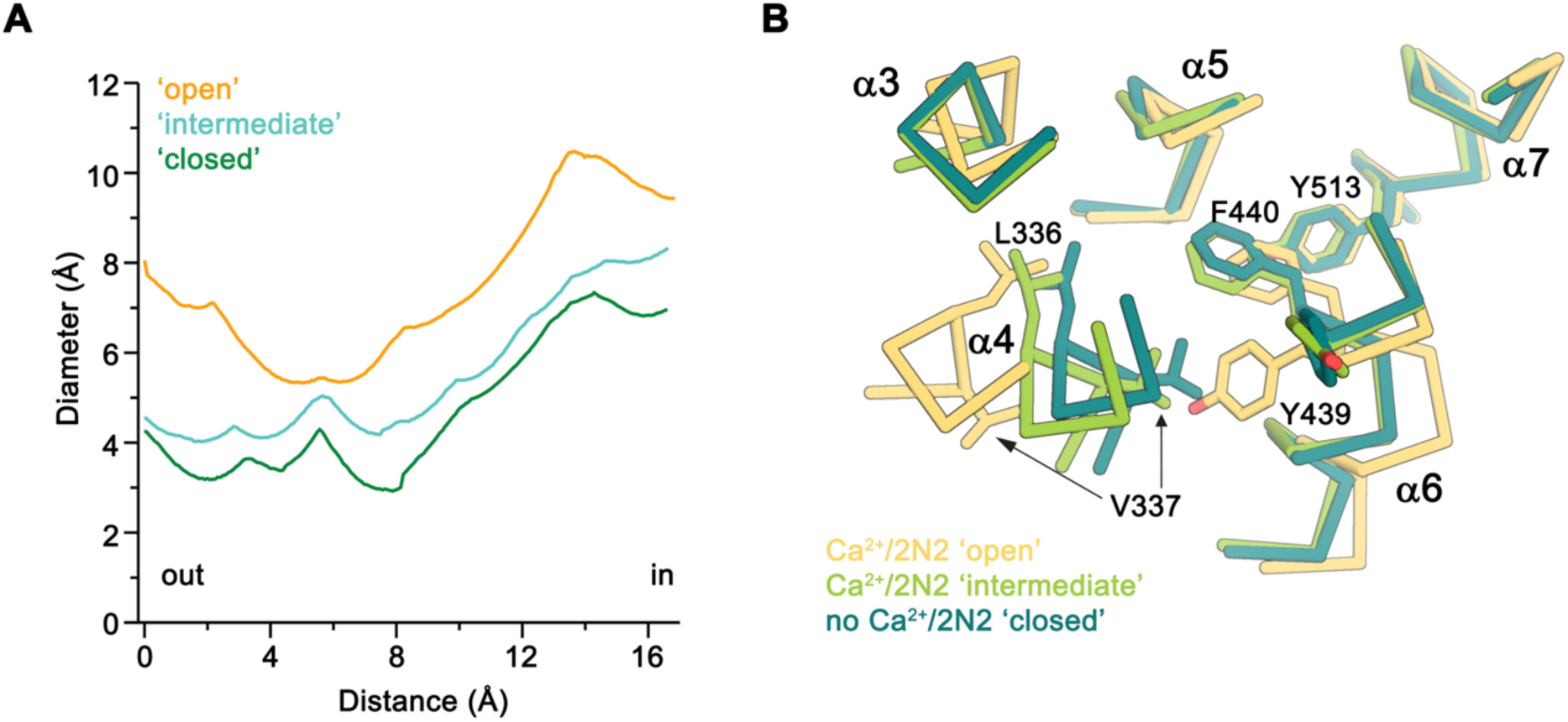
Diameter of the subunit cavity in Ca^2+^-bound nhTMEM16 structures in nanodiscs. (**A**) Pore diameter at the constriction of the subunit cavity in the three conformations of the subunit cavity calculated with HOLE (Smart et al., 1996). (**B**) View of the constriction of the ‘subunit cavity’ in three different conformations from the extracellular side. The protein is shown as Cα-trace with selected sidechains displayed as sticks and labelled.

**Figure 7-figure supplement 1.**
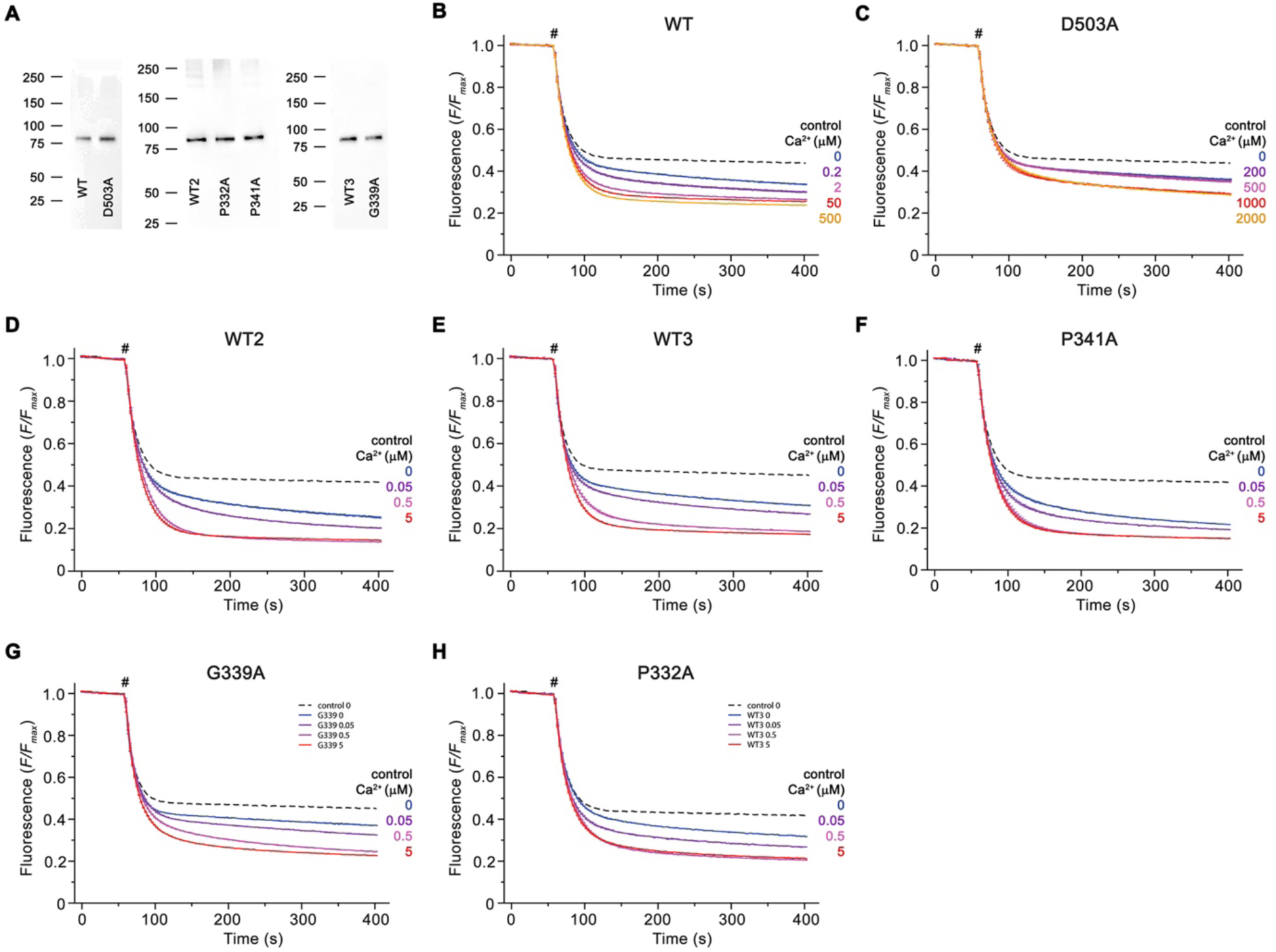
Reconstitution efficiency. (**A**) Reconstitution efficiency of nhTMEM16 mutants compared to WT (WT1, WT2, WT3) reconstituted in the same batch of solubilized lipids. For analysis, equivalent amounts of liposomes were loaded on SDS-PAGE and detected with an anti-my antibody detecting a fusion tag on the protein. The positions of molecular weight markers (kDa) are indicated. (**B**-**H**) Traces of scrambling experiments displayed in Figure 7. Data show averages of three technical replicates, errors are s.e.m.. Ca^2+^-dependence of scrambling activity of WT1 (**B**) and D503A (**C**) displayed in Figure 7B and C. (**D**) Ca^2+^-dependence of scrambling activity of WT2 displayed for comparison in Figures 7D and F. (**E**) Ca^2+^-dependence of scrambling activity of WT3 displayed for comparison in Figure 7E. (**F**) Ca^2+^-dependence of scrambling activity of P341A displayed for comparison in Figure 7D. (**G**) Ca^2+^-dependence of scrambling activity of G339A displayed for comparison in Figure 7E. (**H**) Ca^2+^-dependence of scrambling activity of P332A displayed for comparison in Figure 7F. Traces depict fluorescence decrease of tail-labeled NBD-PE lipids after addition of dithionite (#) at different Ca^2+^ concentrations. Data show averages of three technical replicates. Ca^2+^ concentrations (µM) are indicated.

**Figure 8-figure supplement 1.**
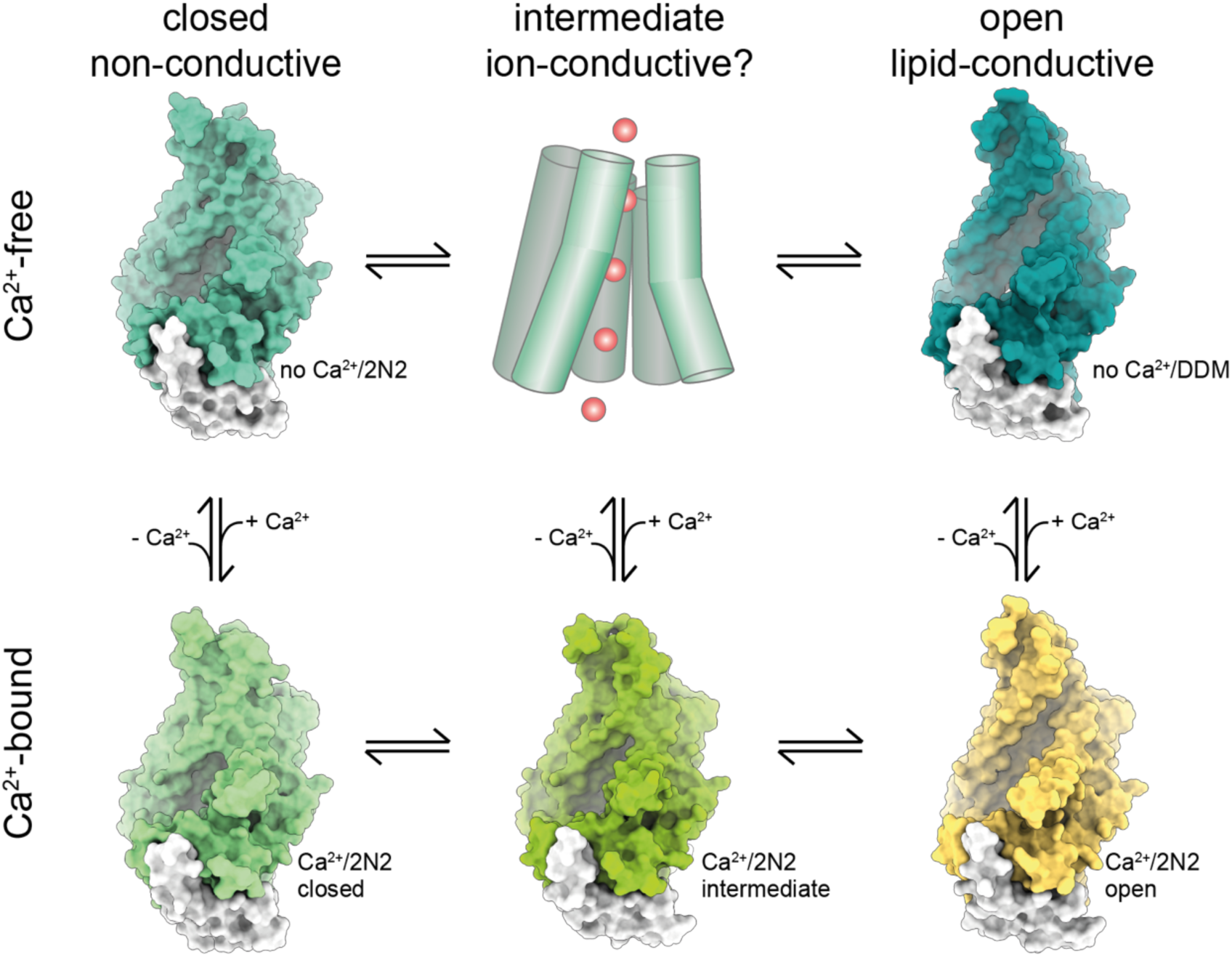
Activation mechanism. Scheme of the stepwise activation of nhTMEM16 as shown in Figure 8 with the states displayed by the surface representation of the respective conformation.

**Video 1. Conformational transitions.** Shown is a morph between distinct conformational states obtained for nhTMEM16, namely: the Ca^2+^-bound ‘open’ state obtained for nhTMEM16 in nanodiscs; the ‘intermediate’ state obtained for nhTMEM16 in lipid nanodiscs; the ‘closed’ state obtained for nhTMEM16 in lipid nanodiscs; and the Ca^2+^-free closed state obtained for nhTMEM16 in lipid nanodiscs. Shown is a view of the subunit cavity as depicted in Figure 3 and 4. Ca^2+^ ions are shown as blue spheres.

**Video 2. Micelle and lipid distortion.** Shown are the cryo-EM density maps of nhTMEM16 obtained in detergent in complex with Ca^2+^ (upper left corner), in detergent in absence of Ca^2+^ (upper right corner), in lipid nanodiscs in complex with Ca^2+^ (lower left corner) and in lipid nanodiscs in absence of Ca^2+^ (lower right corner). The surrounding environment corresponding to the detergent micelle (upper panels) or the lipid nanodiscs (lower panels) are colored respectively, while the rest of the protein is shown in grey. Refined and unmasked cryo-EM maps were low-pass filtered to 6 Å. Transmembrane α-helices 4 and 6 are labeled at the end, indicating the position of the subunit cavity.

